# The Interferon γ Pathway Enhances Pluripotency and X-Chromosome Reactivation in iPSC reprogramming

**DOI:** 10.1101/2023.07.31.551297

**Authors:** Mercedes Barrero, Anna V. López-Rubio, Aleksey Lazarenkov, Enrique Blanco, Moritz Bauer, Luis G. Palma, Anna Bigas, Luciano Di Croce, José Luis Sardina, Bernhard Payer

## Abstract

Reprogramming somatic cells into induced pluripotent stem cells (iPSCs) requires activation of the pluripotency network and resetting of the epigenome by erasing the epigenetic memory of the somatic state. In female mouse cells, a critical epigenetic reprogramming step is the reactivation of the inactive X chromosome. Despite its importance, a systematic understanding of the regulatory networks linking pluripotency and X-reactivation is missing. Here we reveal the pathways important for iPSC reprogramming and X-reactivation using a genome-wide CRISPR screen. In particular, we discover that activation of the interferon γ (IFNγ) pathway early during reprogramming accelerates pluripotency acquisition and X-reactivation. IFNγ stimulates STAT3 signaling and the pluripotency network and leads to enhanced TET-mediated DNA demethylation, which consequently boosts X-reactivation. We therefore gain a mechanistic understanding of the role of IFNγ in reprogramming and X-reactivation and provide a comprehensive resource of the molecular networks involved in these processes.

## Introduction

A characteristic hallmark of embryonic development and pluripotency is extensive epigenetic reprogramming (*1*,*2*), for which the X chromosome is a prime example in female mammals (*3*,*4*). During female mouse development, one of the two X chromosomes switches between active and inactive states in a dynamic fashion, in order to balance gene dosage with autosomes and XY males. The paternally inherited X chromosome is first inactivated during preimplantation development and is then subsequently reactivated in the epiblast of the late blastocyst embryo, the lineage from which all embryonic cell types emerge, and pluripotent embryonic stem cells (ESCs) are derived in culture (*5–7*). The erasure of epigenetic memory during X-chromosome reactivation allows afterwards postimplantation epiblast cells to undergo random X-chromosome inactivation during their exit from naive pluripotency. While X-inactivation is stably maintained in somatic cells, female germ cells go through a second wave of X-chromosome reactivation prior to, and around the time that the cells are entering meiosis and differentiating into oocytes (*8–12*).

Not only during female mouse development *in vivo*, but also in cell culture *in vitro*, the cellular differentiation and X-chromosome states are tightly linked. While differentiated cell types are characterized by X-chromosome inactivation, female mouse pluripotent stem cells such as ESCs and induced-pluripotent stem cells (iPSCs) have two active X chromosomes. On a molecular level, this can be explained by the repressive effect of the pluripotency factor network on the expression of *Xist*, the non-coding master regulator of X-inactivation (*13–16*), coupled with the upregulation of *Xist* activators during differentiation of pluripotent stem cells, thereby triggering random X-chromosome inactivation (*17–21*). X-inactivation in mouse somatic cells is reversed during reprogramming into iPSCs by the process of X-chromosome reactivation (*22*). Previous studies have characterized the kinetics and revealed some of the regulators of X-chromosome reactivation during iPSC reprogramming (*15*, *23–27*), however, the full mechanisms are far from being understood.

The implementation of the CRISPR/Cas9 technology for genome and epigenome editing has allowed the generation of large-scale CRISPR screens based on the expression of pooled gRNA libraries (*28*), leading to the identification of previously unknown players in pluripotency exit (*29*–*32*), maintenance (*33–36*) and acquisition (*37–39*). Furthermore, CRISPR screens in the context of the X chromosome enabled the identification of genes driving sex differences in ESCs (*40*) and *Xist* regulators (*21*, *41*). So far, most perturbation screens on the topic of pluripotency acquisition have relied on the identification of factors constituting roadblocks of the reprogramming process (*39*, *42–46*), as siRNAs, shRNAs or gRNAs targeting those genes would be enriched and therefore easily detected in the final iPSC-population when knocked-out or knocked-down. However, there is a lack of genome-wide screens revealing active players in pluripotency acquisition or X-reactivation, as dropout screens rely on large cell numbers to ensure faithful shRNA/gRNA representation, which has been hard to achieve during iPSC reprogramming due to low reprogramming efficiencies. As a result, only small-scale candidate approaches have been carried out so far to identify drivers of X-chromosome reactivation during somatic cell reprogramming (*15*, *23*, *25*, *27*), and a comprehensive study of the gene regulatory networks controlling this process is missing.

To fill this gap, we performed a genome-wide CRISPR knockout screen during reprogramming of neural precursor cells (NPCs) into iPSCs, with the aim to reveal the pathways important for the process of X-chromosome reactivation. Using this approach, we discovered that the interferon γ pathway regulates reprogramming and X-chromosome reactivation. Our results also show that this process is dependent on TET-mediated DNA demethylation of X-chromosomal genes, thereby facilitating X-reactivation.

## Results

### A genome-wide CRISPR knockout screen reveals molecular networks involved in reprogramming and X-chromosome reactivation

In order to gain a comprehensive understanding of the pathways important for pluripotency acquisition and X-chromosome reactivation, we developed a cell line suitable for a genome-wide CRISPR screen during reprogramming. This approach, based on our PAX (pluripotency and X-chromosome reporter) reprogramming system (*24*), enables us to trace the X-chromosome activity following an X-GFP reporter (see “Materials and Methods”). We further modified this cell line by the introduction of a doxycycline-inducible *Cas9* (*iCas9*) transgene to mediate CRISPR-based target gene deletions (*47*). We then infected these ESCs with a gRNA library targeting all protein-coding genes in the mouse genome (*48*) and differentiated them into neural precursor cells (NPCs) leading to X-chromosome inactivation, as indicated by silencing of the X-GFP reporter (**Fig. 1A**). These NPCs provided the starting material for our reprogramming screen. We then induced reprogramming by adding doxycycline, which activated the expression of an *MKOS* (*c-Myc*, *Klf4*, *Oct4* and *Sox2*) reprogramming cassette (*49*) and the *iCas9* at the same time, resulting in the production of knockouts during the reprogramming process. After ten days of reprogramming, we used fluorescence-activated cell sorting (FACS) to separate three different populations, based on the SSEA1 pluripotency marker and X-GFP (indicative of a reactivated X chromosome). The three isolated populations were classified as non-pluripotent (SSEA1-X-GFP-), early pluripotent (SSEA1+ X-GFP-) and late pluripotent (SSEA1+ X-GFP+). By comparing the abundance of gRNAs and their enrichment or depletion across different populations we finally identified genes with different roles for the reprogramming and X-chromosome reactivation processes.

**Fig. 1.**
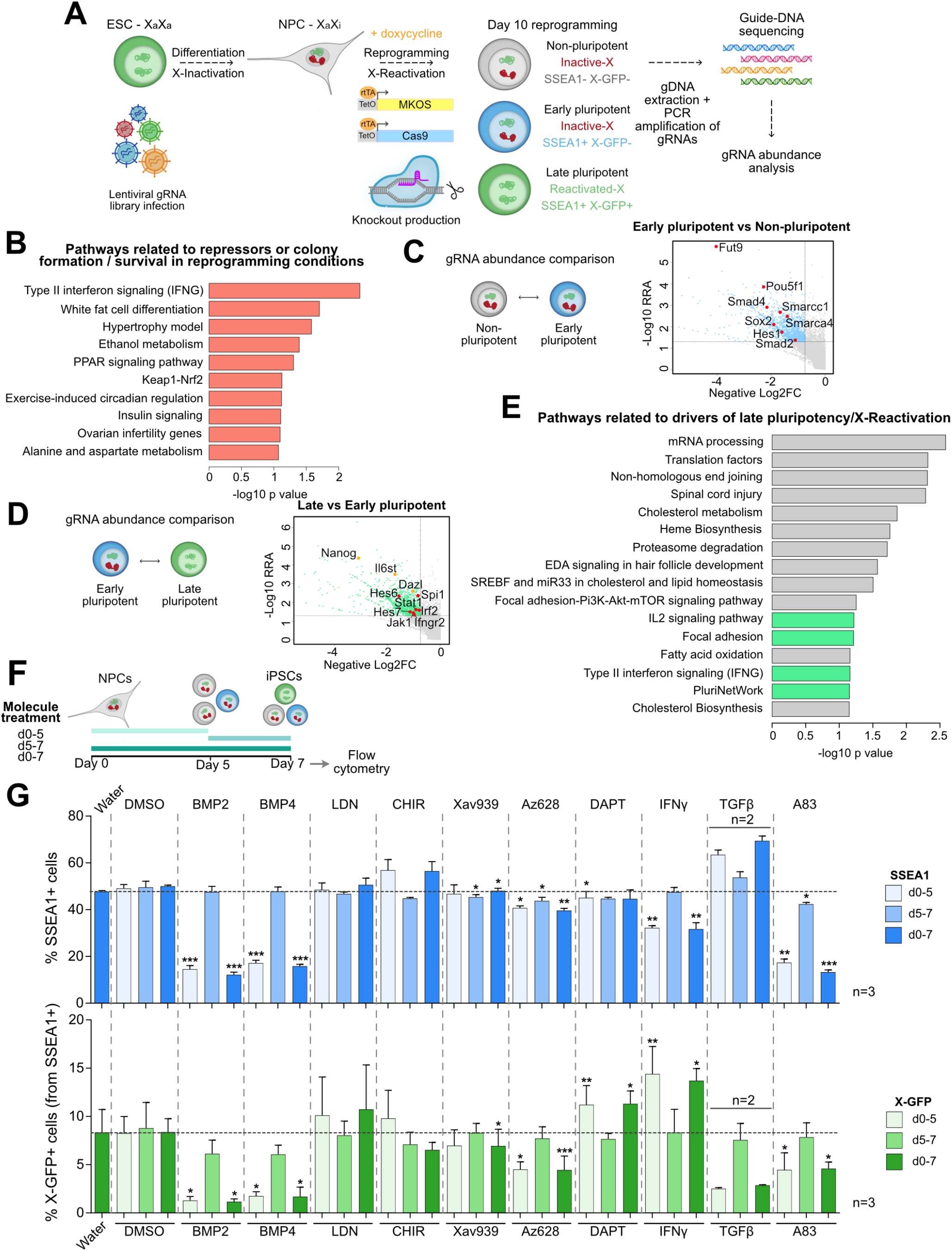
A genome-wide CRISPR knockout screen reveals molecular networks involved in reprogramming and X-chromosome reactivation. (**A**) Experimental design. X-GFP *iCas9* ESCs were infected with a genome-wide lentiviral gRNA library and differentiated into NPCs. Next, NPCs were treated with doxycycline to activate the expression of the reprogramming cassette and the *iCas9*. The knockouts took place during reprogramming. At day 10 of reprogramming, three populations were FACS-separated: non-pluripotent (SSEA1-X-GFP-), early pluripotent (SSEA1+ X-GFP-) and late pluripotent, X-reactivated (SSEA1+ X-GFP+). For these three populations and the NPCs, genomic DNA extraction, PCR-amplification of the gRNA sequences, sequencing and analysis of gRNA abundance were performed. n=2 biological replicates (independent reprogramming rounds). (**B**) Pathways related to overrepresented genes in the three reprogramming populations (non-pluripotent, early pluripotent and late pluripotent) compared to NPCs (WikiPathways Mouse 2019). For pathway enrichment analysis, the top 250 genes from each individual comparison to NPCs ranked by RRA score with MAGeCK software were merged and selected. (**C**) gRNA abundance comparison (early pluripotent vs non-pluripotent) and representation of genes with negative Log2FC (underrepresented) vs −log10 RRA (RRA cutoff = 0.05, Log2FC cutoff = −0.75) (activators of early pluripotency). (**D**) gRNA abundance comparison (late pluripotent vs early pluripotent) and representation of genes with negative Log2FC (underrepresented) vs −log10 RRA (RRA cutoff = 0.05, Log2FC cutoff = −0.75) (activators of late pluripotency, X-reactivation). Genes highlighted in yellow are related to pluripotency, genes highlighted in red are involved in Notch or interferon γ signaling. (**E**) Pathways (WikiPathways Mouse 2019) related to underrepresented genes in “late pluripotent vs early pluripotent” comparison (activators of late pluripotency, X-reactivation, n=1313 genes). Pathways related to proliferation, differentiation and metabolism are shown in gray. The rest of the pathways are highlighted in green. RRA score < 0.05 and Log2FC < −0.8 filtering was applied. (**F**) Experimental design for (G). Treatment with molecules targeting pathways identified in the CRISPR screen was done at the beginning of reprogramming (day 0 - day 5), later in reprogramming (day 5 - day 7) or during the whole process (day 0 - day 7). At day 7, flow cytometry analysis of SSEA1 and X-GFP percentages was performed. (**G**) Pathway validation by molecule treatment at the beginning of reprogramming (day 0 - day 5), later in reprogramming (day 5 - day 7) or during the whole process (day 0 - day 7) in three independent reprogramming rounds, n=3 (except for TGFβ, n=2). Flow cytometry analysis at day 7 of SSEA1 and X-GFP percentages is shown. Data represented as mean +/− SD. Statistics (paired t-tests): where not specified = non-significant; * = p<0.05; ** = p<0.01; *** = p<0.001.

Genes required for cell survival and normal growth (the essentialome) were depleted in all three final cell populations (non-pluripotent, early pluripotent and late pluripotent) when compared to NPCs (**Fig. S1C-G**). On the other hand, overrepresented genes in the three reprogramming populations constituted repressors of iPSC-colony formation and cell survival in reprogramming conditions. We found enrichment of pathways related to differentiation, metabolism and inflammation (**Fig. 1B**). “Type II Interferon signaling” / Interferon γ (IFNγ) pathway showed the highest overrepresentation, suggesting a putative role of this pathway in repressing colony formation during reprogramming.

Next, in order to identify genes and pathways with a role early during pluripotency acquisition, we compared gRNA frequencies between the non-pluripotent and early pluripotent populations (**Fig. 1C**, **Fig. S1H-J**). As expected, we found genes with well-known roles in pluripotency such as *Smad2*, *Smad4* (related to BMP signaling), *Pou5f1*, *Sox2*, *Smarcc2*, *Smarca4* (related to pluripotency), *Hes1* (target of Notch pathway) and *Fut9* (that encodes the key enzyme necessary for SSEA1 synthesis), thereby validating our screening approach (**Fig. 1C**).

As our main aim was to identify new genes and pathways playing a role in naive pluripotency and in particular X-chromosome reactivation, we then focused on the comparison between the late and early pluripotent populations (**Fig. 1D and 1E**, **Fig. S1K and S1L**). Among the genes and pathways identified as drivers of naive pluripotency and X-reactivation, we found processes involved in cell proliferation (mRNA processing, translation), lipid metabolism, the pluripotency network (with genes like *Nanog*, *Il6st* and *Dazl*), the Notch pathway (*Hes6* and *Hes7*) and the interferon γ pathway (including genes such as *Stat1*, *Jak1*, *Spi1*, *Irf2* and *Ifngr2*) (**Fig. 1D and 1E**).

Next, we validated our screening results by activating and/or repressing some of the identified pathways through the addition of signaling factors and small molecules during the reprogramming process focusing on potential regulators of colony formation, pluripotency acquisition or X-chromosome reactivation. In order to identify an early or late contribution of the different pathways, treatment was performed at the beginning of reprogramming (from day 0 to 5), at the end of reprogramming (from day 5 onwards), or during the whole process (**Fig. 1F**). We tested the following pathways: BMP (activated by BMP2 and BMP4, repressed by LDN-212854), Wnt (activated by the GSK-3β inhibitor CHIR99021, repressed by the tankyrase1/2 inhibitor Xav939), MAPK (repressed by the pan-Raf kinase inhibitor Az628), Notch (inhibited by the γ-secretase inhibitor DAPT), interferon γ pathway (activated by IFNγ), and TGFβ (activated by TGFβ, repressed by the ALK5, ALK4 and ALK7 selective inhibitor A83-01). We measured the effects of the treatments on early pluripotency (SSEA1+) and X-chromosome reactivation (X-GFP+) by flow cytometry on day 7 of reprogramming, when we observed the onset of X-chromosome reactivation, and therefore, the most dramatic change in X-chromosome status (from inactive to active) (**Fig. 1G**). Some molecules, such as BMP2, BMP4, or A83-01, caused a reduction of both SSEA1 and X-GFP percentages upon early or continuous treatment, indicating an early effect in the process of reprogramming. By contrast, the early or continuous treatment with IFNγ (activator of IFNγ pathway) and DAPT (inhibitor of Notch pathway) resulted in an increased percentage of X-GFP+ cells (around 1.7 and 1.3 fold, respectively) without increasing the percentage of SSEA1+ cells. This suggests a putative role of these molecules in the later stages of reprogramming.

Overall, our CRISPR screen identified both known and novel pathways related to the different reprogramming stages. Interestingly, we found interferon γ signaling to play contrasting roles during the iPSC-reprogramming process: early on as a repressor of colony formation (**Fig. 1B**), but subsequently as a driver of late pluripotency and X-chromosome reactivation (**Fig. 1D and 1E**). As IFNγ induced the highest increase in X-chromosome reactivation efficiency in the validation experiments and has never been implicated in these processes before, we therefore from now on focused on characterizing its mechanism of action.

### Interferon γ signaling modulates colony formation and X-chromosome reactivation during iPSC reprogramming

In our CRISPR screen, the IFNγ pathway showed up as a putative repressor of iPSC-colony formation and potential driver of X-chromosome reactivation. We explored the role of IFNγ signaling in these two scenarios after IFNγ treatment at different time points: early (day 0-5), late (day 5-10) and continuous (day 0-10) (**Fig. 2A**). To further investigate the implication of the IFNγ pathway activation in iPSC colony formation, we performed Alkaline Phosphatase (AP) staining after 10 days of reprogramming (**Fig. 2B**) upon different timings of IFNγ treatment. The early (day 0-5) and continuous (day 0-10) treatments induced a statistically significant decrease in AP-positive colony number, validating the role of IFNγ signaling as a repressor of colony formation, while the late treatment (day 5-10) did not have any effect. This phenotype could be related to a slight increase in apoptosis observed after 48h from IFNγ treatment during the onset of reprogramming induction (**Fig. S2A**). Next, we tested X-chromosome reactivation efficiency by measuring the percentage of X-GFP positive cells at days 5, 7 and 10 of reprogramming (**Fig. 2C-E**). At day 7, the early and continuous treatment with IFNγ resulted in a significant increase in cells undergoing X-GFP reactivation (**Fig. 2D**), with average fold changes of 1.75 and 1.71 to the control, respectively (**Fig. 2E**), while the differences between IFNγ-treated and control samples were less prominent at day 10 (**Fig. 2D and 2E**), suggesting that early IFNγ treatment accelerates X-reactivation.

**Fig. 2.**
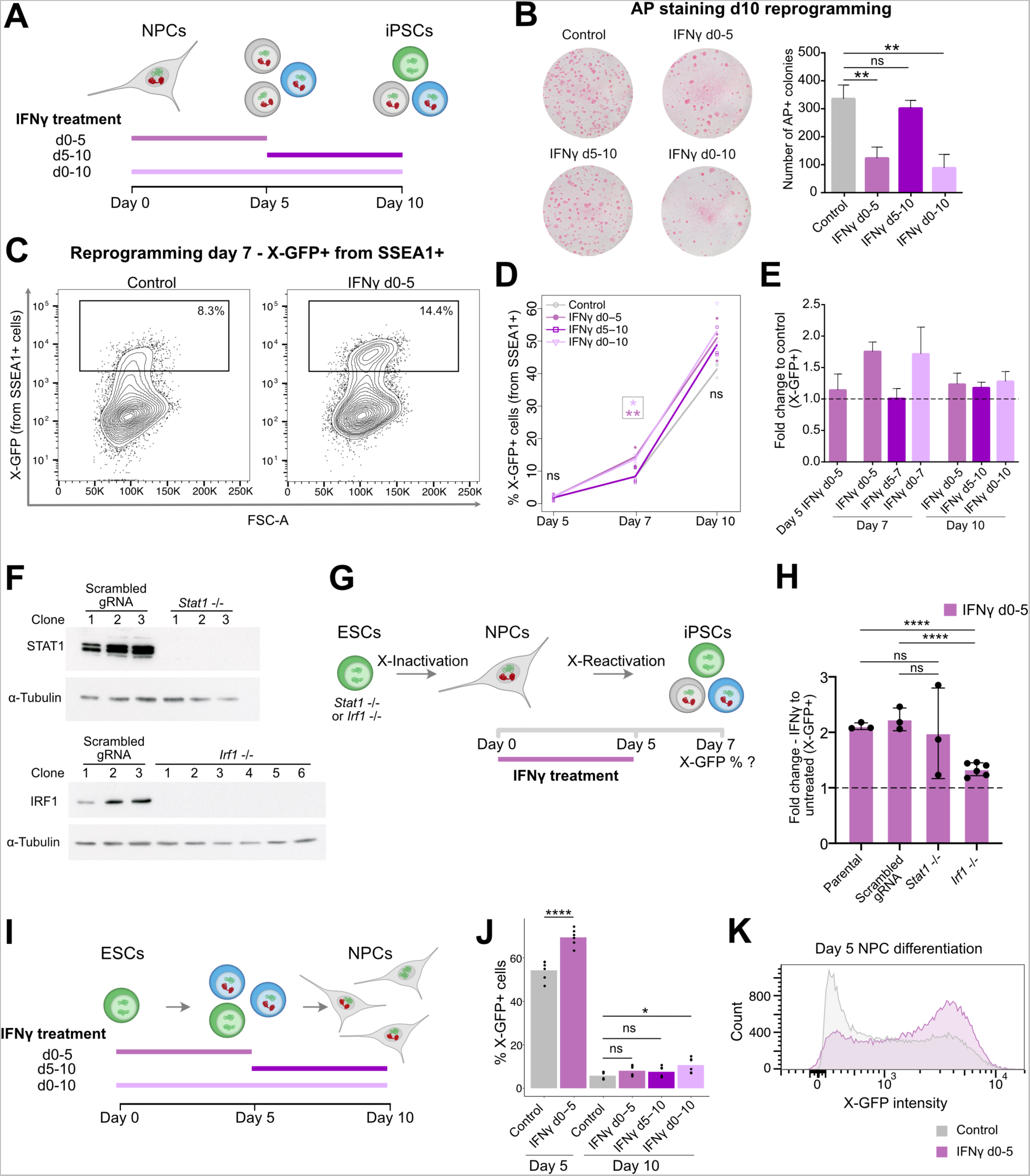
Interferon γ signaling modulates colony formation and X-chromosome reactivation during iPSC reprogramming. (**A**) Experimental design for (B-E): IFNγ treatment was done at the beginning of reprogramming (day 0 - day 5), at the end of reprogramming (day 5 - day 10) or during the whole process (day 0 - day 10) in three independent reprogramming rounds, n=3. (**B**) Alkaline Phosphatase (AP) stainings on day 10 of reprogramming in control and IFNγ treatments (d0-5, d5-10, d0-10) and counting of AP+ colonies (n=3 for control, n=4 for IFNγ treatments). Statistics (unpaired t-tests): ns = non-significant; ** = p<0.01. Error bars represent SD. (**C**) Flow cytometry plots of X-GFP expression (from SSEA1+ cells) in control and IFNγ treatment (day 0-5) on day 7 iPSCs. Gating shows the X-GFP+ population. (**D**) Flow cytometry analysis of X-GFP+ cells (from SSEA1+) on days 5, 7 and 10 of reprogramming, in control and IFNγ treatments (day 0-5, day 5-10, day 0-10). Statistics (paired t-tests): where not specified = non-significant; * = p<0.05; ** = p<0.01. (**E**) Fold change of X-GFP percentages (from SSEA1+ cells) in IFNγ treatments compared to controls, on days 5, 7 and 10 of reprogramming, measured by flow cytometry. Error bars represent SD. Calculations are done based on percentages shown in (D). (**F**) Western blot of STAT1 for three scrambled-gRNA clones and 3 *Stat1 −/−* clones (top) and IRF1 for three scrambled-gRNA clones and 6 *Irf1 −/−* clones (bottom). ɑ-Tubulin was used as loading control. (**G**) Experimental design for (H): *Stat1−/−*, *Irf1−/−*, parental and scrambled gRNA control ESCs were differentiated into NPCs and then reprogrammed into iPSCs in the presence or absence of IFNγ (day 0-5). X-GFP percentages (from SSEA1+ cells) were measured by flow cytometry at day 7 of reprogramming. 3 clones from the parental cell line, 3 clones containing a scrambled gRNA, 3 *Stat1 −/−* clones and 6 *Irf1 −/−* clones were used, including three technical replicates for each clone. (**H**) Fold change of percentage of X-GFP+ cells (from SSEA1+ cells) in IFNγ-treated cells compared to untreated controls on day 7 of reprogramming, measured by flow cytometry. Bars represent the average X-GFP fold change (IFNγ vs control) for clones with the same genotype, listed in (G). Each dot represents the mean of three technical replicates for each clone. Statistics (unpaired t-tests): ns = non-significant; **** = p<0.0001. Error bars represent SD. (**I**) Experimental design for (J-K): NPC differentiation was induced from ESCs, and treatment with IFNγ was done from day 0 to 5, from day 5 to 10, or from day 0 to 10 of differentiation (n=6 independent replicates). (**J**) Quantification of X-GFP percentage on days 5 and 10 of NPC differentiation by flow cytometry in control and IFNγ treatment conditions. Statistics (paired t-tests): ns = non-significant; *=p<0.05; **** = p<0.0001. (**K**) Flow cytometry histogram of X-GFP intensity in representative samples of control and IFNγ-treated day 5 NPCs.

IFNγ signaling induces the activation of the transcription factors STAT1 and IRF1, which in turn activate the expression of IFNγ-response genes. In order to determine the speed of activation of interferon γ target genes upon treatment, we analyzed the expression of *Irf1* and *Gbp2* by RT-qPCR in NPCs during the first 9 hours of reprogramming induction. A strong increase in the expression of the IFNγ pathway genes *Irf1* and *Gbp2* was observed already after 3-6 hours of treatment (**Fig. S2B**). Moreover, an increased expression of STAT1 and phospho-STAT1 at the protein level was observed at days 2 and 5 of reprogramming in the IFNγ-treated cells compared to the control, indicating activation of the pathway during reprogramming upon IFNγ treatment (**Fig. S2C-E**). To shed light on the mechanism behind the increased X-chromosome reactivation efficiency upon IFNγ treatment, we generated *Stat1*−/− and *Irf1*−/− ESC lines (**Fig. 2F**), induced reprogramming in NPCs generated from them with and without IFNγ treatment from day 0 to 5, and analyzed the percentages of cells undergoing X-GFP reactivation at day 7 of reprogramming by flow cytometry (**Fig. 2G and 2H**, **Fig. S2F**). As in our previous experiments, IFNγ treatment resulted in an around 2-fold increase in the percentage of X-GFP+ cells in the parental and scrambled gRNA controls compared to untreated cells. In the *Stat1*−/− cell lines, IFNγ treatment still induced an increase in X-GFP reactivation efficiency comparable to the controls in two out of three clones (about 2-fold), suggesting that STAT1 is unlikely to be the main responsible downstream factor for the observed phenotype. By contrast, all six *Irf1*−/− clones analyzed showed a reduced increase in X-GFP reactivation compared to the untreated control clones (IFNγ vs control X-GFP fold changes varied from 1.18 to 1.46, p<0.0001). Together, these data suggest that IRF1, but not STAT1, is a mediator of IFNγ signaling responsible for the increased efficiency of X-GFP reactivation observed upon IFNγ treatment.

Next, we explored if IFNγ treatment has the opposite effect on ESC differentiation into NPCs. For this, we treated cells undergoing differentiation with IFNγ from day 0 to 5, day 5 to 10 or throughout the whole process (**Fig. 2I**) and assessed the percentages of SSEA1+ and X-GFP+ cells by flow cytometry on days 5 and 10 (**Fig. S2G**, **Fig. 2J and 2K**). At day 5 of differentiation, no changes in SSEA1 percentage were detected between the control and the IFNγ-treated samples (**Fig. S2G**), while the X-GFP percentage was elevated in the IFNγ-treated cells at day 5 of differentiation compared to the control (**Fig. 2J and 2K**). In contrast, on day 10 of differentiation, substantial changes were no longer detected in SSEA1 or X-GFP expression between control and treated samples (**Fig. S2G**, **Fig. 2J and 2K**). These data indicate that IFNγ treatment during differentiation delays X-chromosome inactivation, which is opposite to its observed role in NPC to iPSC reprogramming, where IFNγ accelerates X-chromosome reactivation instead.

### Interferon γ pathway activation accelerates the reprogramming process

In order to gain insight into the changes induced by early IFNγ treatment (day 0-5), we performed transcriptomic analyses of FACS-sorted cells at days 2, 5 (SSEA1+) and 7 (SSEA1+/X-GFP negative, medium and high; **Fig. S3A**) of reprogramming and compared them to untreated cells, and NPCs and ESCs as fully differentiated and pluripotent cell types, respectively. Principal Component Analysis (PCA) revealed a strong similarity between control and IFNγ-treated iPSCs at days 2 and 5 (**Fig. 3A**). However, at day 7, IFNγ-treated iPSCs showed an accelerated reprogramming kinetics compared to the control, clustering closer to the ESCs.

**Fig. 3.**
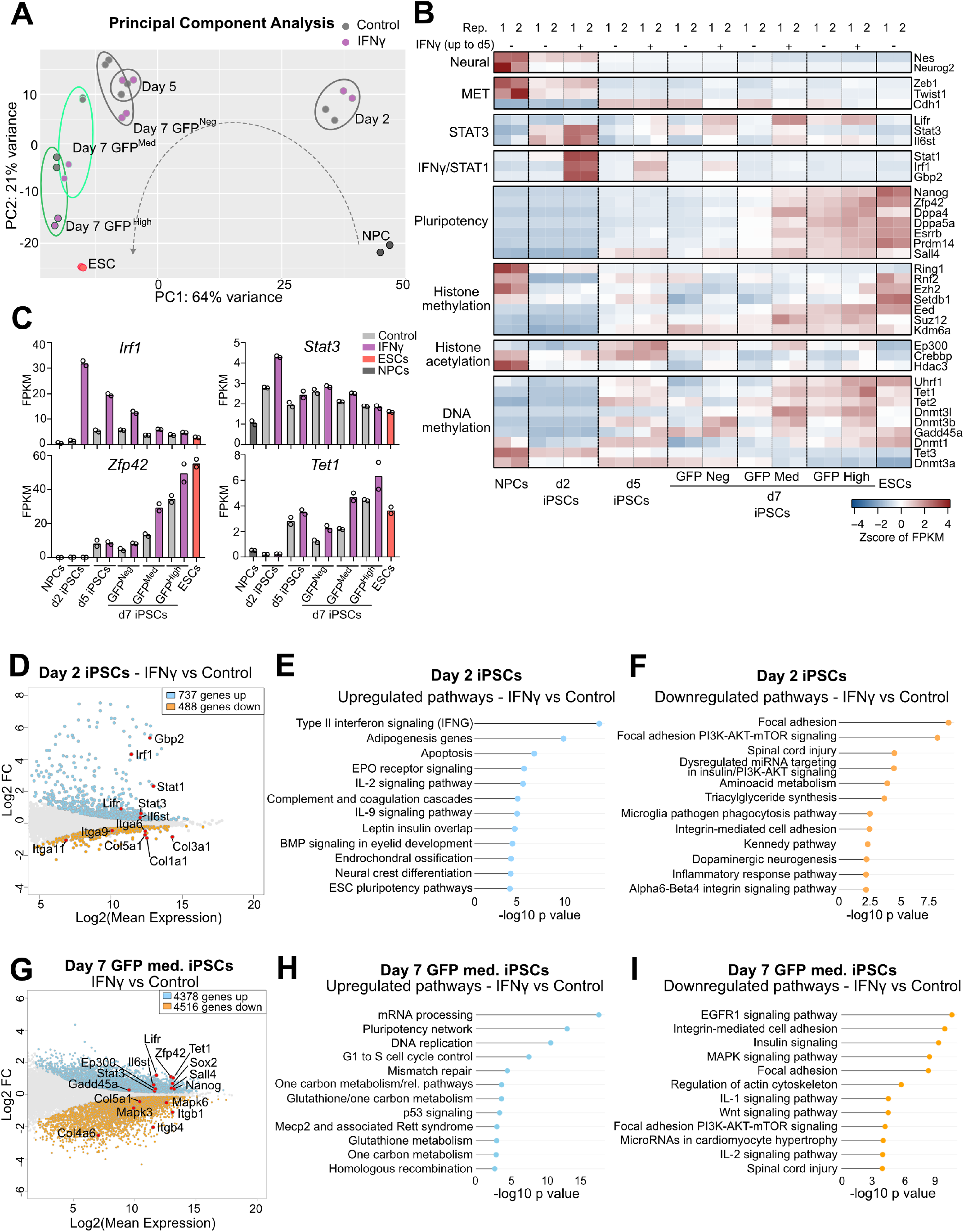
Interferon γ pathway activation accelerates the reprogramming process. (**A**) Principal component analysis of RNA-sequencing of NPCs, day 2, day 5, day 7 iPSC populations and ESCs, in control and IFNγ treatment (day 0-5), representing the top 500 most variable genes. (**B**) Heatmap representing expression (Z score of FPKM) of neural genes, mesenchymal-to-epithelial transition (MET) genes, pluripotency genes, STAT3- and IFNγ/STAT1-related genes, histone methylation genes, histone acetylation genes and DNA methylation genes. (**C**) Expression (FPKM) of selected genes (*Irf1, Stat3, Zfp42/Rex1* and *Tet1*) in NPCs, ESCs, day 2, day 5 and day 7 iPSC populations +/−IFNγ treatment (two RNA-sequencing replicates shown). (**D**) MA plot displaying transcriptomic changes of IFNγ vs control d2 iPSCs (adjusted p value = 0.1). Upregulated genes are highlighted in light blue, downregulated genes are highlighted in orange. Selected genes are shown with points in red. (**E, F**) Upregulated (E) and downregulated (F) pathways of IFNγ vs control d2 iPSCs (WikiPathways Mouse 2019) (adjusted p value = 0.1). (**G**) MA plot displaying transcriptomic changes of IFNγ vs control d7 X-GFP medium iPSCs (adjusted p value = 0.1). Upregulated genes are highlighted in light blue, downregulated genes are highlighted in orange. Selected genes are shown with points in red. (**H, I**) Upregulated (H) and downregulated (I) pathways of IFNγ vs control d7 X-GFP medium iPSCs (WikiPathways Mouse 2019) (adjusted p value = 0.1).

Next, we explored the expression of several genes involved in the reprogramming process (**Fig. 3B and 3C, Fig. S3B**). This analysis indicated an early activation of the IFNγ-related genes *Stat1*, *Irf1* and *Gbp2* upon treatment, which peaked at day 2 of reprogramming. Of note, IFNγ-treated iPSCs also showed a higher expression of some genes related to the LIF/STAT3 pathway (*Stat3*, *Lifr*, *Il6st*), which is involved in the acquisition of pluripotency (*50–52*). Importantly, pluripotency genes showed a significantly higher expression in the IFNγ-treated samples compared to the control, especially in the X-GFP-medium population, which is undergoing X-reactivation. Examples of these naive pluripotency genes are *Nanog*, *Zfp42/Rex1*, *Dppa4*, *Dppa5a*, *Esrrb*, *Prdm14* and *Sall4.* This supports a more advanced reprogramming in the IFNγ-treated samples, consistent with what we have observed in the PCA (**Fig. 3A**) and is in line with studies showing an involvement of naive pluripotency factors such as *Nanog* and *Prdm14* in X-chromosome reactivation (*10*, *15*, *53*). We then focused on genes related to DNA demethylation dynamics, as demethylation of X-linked gene promoters is a key step in X-chromosome reactivation (*23*, *25*, *54*). While we did not observe differences in the expression of *Tet2* and *Tet3*, we saw a higher expression of *Tet1* and *Gadd45a* from day 5 onwards in the IFNγ-treated cells in comparison to the control (**Fig. 3B and 3C**). *Tet1* has previously been shown to be upregulated during iPSC reprogramming, and to demethylate and reactivate pluripotency genes (*55*). GADD45A is a member of the base excision repair pathway that was found to interact with TET1, promoting its activity and enhancing DNA demethylation (*56*). Thus, the upregulation of these genes upon IFNγ treatment could induce DNA demethylation leading to a more rapid cell fate transition and more efficient or faster X-chromosome reactivation.

We then performed differential expression analysis between control and IFNγ-treated day 2, day 5 and day 7 iPSCs (**Fig. 3D-I**, **Fig. S3C-H**). As expected, day 2 and 5 iPSCs showed an upregulation of interferon γ signaling pathway signature genes, including *Gbp2*, *Stat1* and *Irf1* in the IFNγ-treated cells (**Fig. 3D and 3E, Fig. S3C and S3D**). Additionally, at day 2 we observed an activation of other inflammation pathways, like complement and coagulation cascades, IL-2, IL-9, and also apoptosis (**Fig. 3E**), fitting with the increased percentage of annexin V-positive cells observed upon IFNγ treatment early during reprogramming (**Fig. S2A**). As mentioned above, some genes from the pluripotency-related STAT3 pathway showed an increased expression early upon interferon γ treatment, like *Lifr*, *Stat3* and *Il6st* (**Fig. 3B-D**), in line with the higher expression of genes related to pluripotency and/or DNA demethylation detected at day 5 (*Esrrb*, *Lifr*, *Tet1* and *Gadd45a*) (**Fig. 3B and 3C**, **Fig. S3B-D**). Focusing on the downregulated pathways and genes upon interferon γ treatment, we found a reduction of focal adhesion genes on both days 2 and 5 (**Fig. 3F and S3E**), predominantly represented by integrins and collagens (*Itga9*, *Col1a1*, *Col3a1*, *Col5a1*) (**Fig. 3D**, **Fig. S3C**). Integrin-mediated cell adhesion has been shown to have an impact in colony number in reprogramming (*43*). Thus, the decreased expression of focal adhesion genes, together with the increased apoptosis observed upon IFNγ treatment (**Fig. S2A**), could explain the lower colony number in the IFNγ-treated samples (**Fig. 2B**).

Next, we compared the transcriptome of day 7 IFNγ-treated X-GFP-negative, X-GFP-medium and X-GFP-high populations with their respective untreated controls. Pairwise comparisons between these populations showed very similar results (**Fig. 3G-I**, **Fig. S3F-H**). In all cases, the early treatment with IFNγ showed an upregulation of proliferation pathways (mRNA processing, G1 to S cell cycle control), metabolism-related pathways, and importantly, the pluripotency network, including genes such as *Nanog* and *Zfp42/Rex1* (**Fig. 3G and 3H**, **Fig. S3F and S3G**). Other genes found to be upregulated in the IFNγ-treated iPSCs were the genes involved in DNA demethylation *Tet1* and *Gadd45a* (**Fig. 3G**, **Fig. S3F**), as also observed at day 5 (**Fig. S3C**). In addition, in the X-GFP-negative and X-GFP-medium populations, several genes belonging to the LIF-STAT3 pathway were found to be upregulated, such as *Il6st*, *Lifr* and *Stat3* (**Fig. 3G**, **Fig. S3F**), consistent with the results of day 2 (**Fig. 3D**). Among the common downregulated pathways in the IFNγ-treated day 7 iPSCs, we found the EGFR1 signaling and MAPK pathways (that are linked to differentiation) (*57*, *58*), inflammation pathways (IL1 and IL2) and also focal adhesion (**Fig. 3I**, **Fig. S3H**), consistent with our previous results on day 2 and day 5. Overall, our transcriptomic analysis revealed that IFNγ-early treatment accelerated the reprogramming process, as reflected by increased expression of STAT3-, DNA demethylation- and pluripotency-related genes.

### Interferon γ treatment during reprogramming increases the activation of JAK-STAT3 signaling, the expression of pluripotency genes and enhances X-chromosome reactivation

To explore if the increased expression of LIF-STAT3 signaling-related genes (**Fig. 3B-D**) correlated with a higher activation of the pathway, we determined the levels of phosphorylated (Tyr705) STAT3 protein by immunofluorescence in control and IFNγ-treated day 2 iPSCs (**Fig. 4A**). Albeit we observed nuclear staining of phospho-STAT3 in the control samples, the signal was more intense in the IFNγ-treated cells, indicating a higher activation of the pathway upon IFNγ treatment. We confirmed this quantitatively by western blot, which showed around 3-fold increased levels of both total and phospho-STAT3 in the IFNγ-treated day 2 iPSCs compared to the control (**Fig. 4B**). However, this effect was no longer observed in IFNγ-treated day 5 iPSCs (**Fig. 4B**), indicating that IFNγ-mediated increase of JAK-STAT3 signaling activation occurs only transiently early during reprogramming. To put this in context with the reprogramming speed, we next calculated a pluripotency score based on the mRNA expression of selected naive pluripotency genes for each time point in control and IFNγ-treated iPSCs (**Fig. 4C**). This score was higher in all day 7 IFNγ-treated iPSCs (X-GFP negative, medium and high) compared to their control counterparts.

**Fig. 4.**
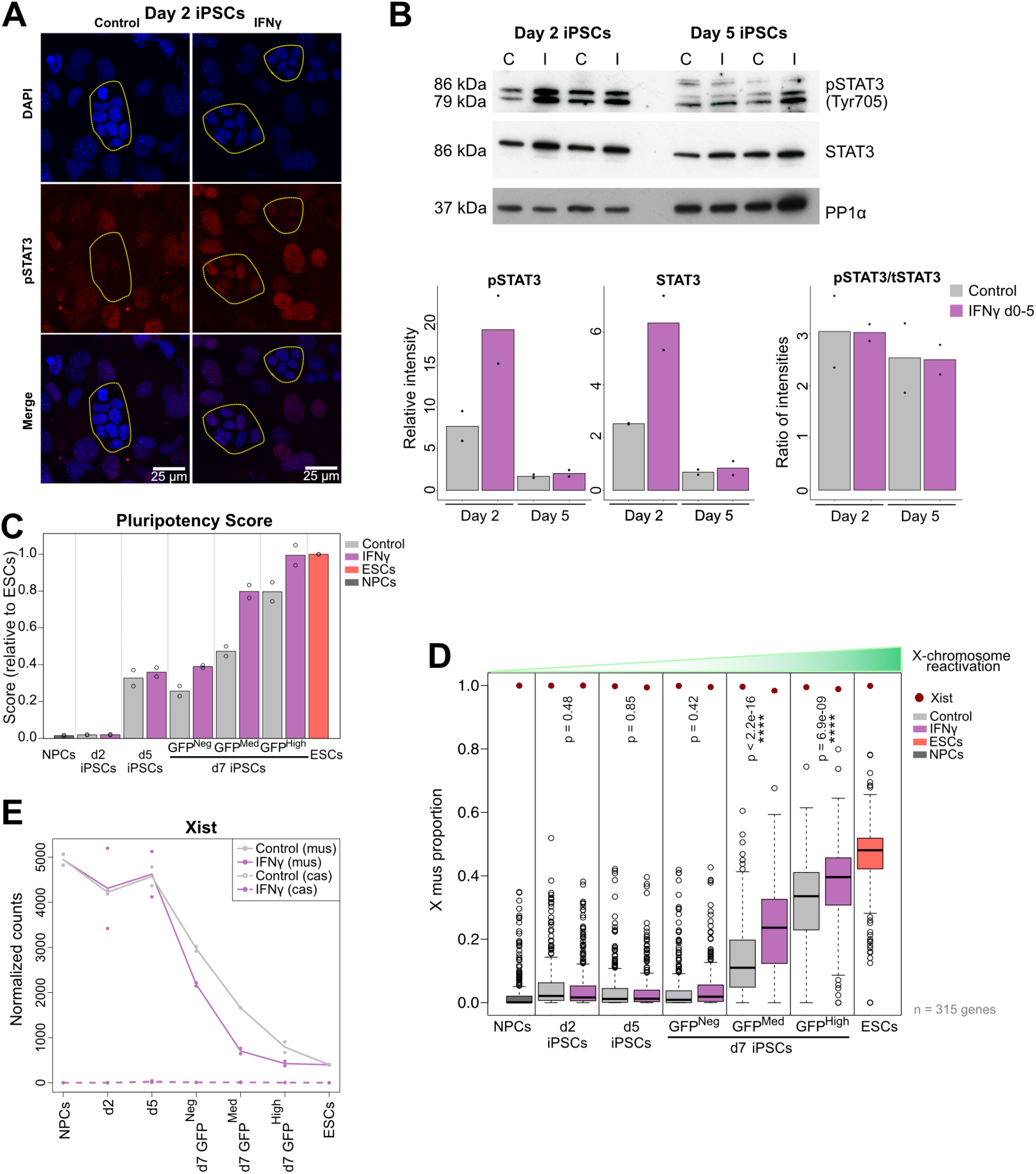
Interferon γ treatment during reprogramming increases the activation of JAK-STAT3 signaling, the expression of pluripotency genes and enhances X-chromosome reactivation. (**A**) Immunofluorescence of pSTAT3 (Tyr705) on day 2 iPSCs +/− IFNγ treatment. Scale bar = 25 µm. Z projections of maximum intensity from 6 stacks are shown for all channels. Outlines highlight colonies of cells undergoing reprogramming, characterized by smaller nuclei and tight aggregation. (**B**) Western blotting of STAT3 and pSTAT3 (Tyr705) on day 2 and day 5 iPSCs +/− IFNγ treatment (C=control, I= IFNγ, loading control: PP1α). Relative intensities to loading control are shown for pSTAT3 and total STAT3, as well as ratio of intensities (phospho-STAT3/total-STAT3). (**C**) Pluripotency score (relative to ESCs) in the two RNA-sequencing replicates during reprogramming, calculated from the expression levels of *Nanog*, *Zfp42/Rex1*, *Dppa4*, *Dppa5a*, *Esrrb*, *Prdm14* and *Sall4*. (**D**) Allelic ratio (X mus proportion) of 315 genes that showed over 25% of total X-linked gene expression in the X cas on NPCs, ESCs, day 2, day 5 and day 7 iPSC populations +/− IFNγ treatment. Statistics: unpaired t-tests. (**E**) Expression (normalized counts) of *Xist* from X mus and X cas in NPCs, ESCs, day 2, day 5 and day 7 iPSC populations +/− IFNγ treatment (two RNA-sequencing replicates shown).

As pluripotency acquisition is linked to X-chromosome reactivation during reprogramming, we analyzed the level of X-chromosome reactivation based on the allelic ratio of X-linked gene expression in each of the iPSC samples. For this, we calculated the X mus proportion in NPCs, day 2, day 5, day 7 iPSCs and ESCs (**Fig. 4D**). When comparing the allelic ratio of IFNγ-treated cells to their control counterparts, we observed a significantly increased X mus proportion in IFNγ-treated iPSCs on day 7, when they undergo X-GFP reactivation. These results showed that not only X-GFP reactivation is more efficient (**Fig. 2C-E**), but also endogenous chromosome-wide X-linked gene reactivation is more advanced upon early activation of the interferon γ pathway. Then, we analyzed the expression of genes from the X-inactivation center, a complex locus containing several coding and non-coding genes that control the expression of *Xist*, the master regulator of X-chromosome inactivation (*59*) (**Fig. 4E**, **Fig. S4A**). We observed that *Xist* expression from the X mus chromosome was consistently lower in the IFNγ-treated cells in the X-GFP-negative, medium and high populations at day 7, in comparison to the control (**Fig. 4E**), while the expression of *Xist* regulators at the X-inactivation center did not show clear changes (**Fig. S4A**). Therefore, it is likely that the accelerated expression of naive pluripotency genes such as *Prdm14* and *Nanog* (**Fig. 3B, Fig. S3B and S4B-E**), which are known to repress *Xist* (*13*, *15*), contribute to the more efficient and advanced X-chromosome reactivation induced by IFNγ, rather than the *Xist* regulators at the X-inactivation center. Interestingly, *Xist* downregulation in the IFNγ-treated day 7 X-GFP-negative iPSCs (**Fig. 4E**) was not sufficient to induce a higher X mus proportion in this cell population (**Fig. 4D**). This could either be due to the not yet complete *Xist* downregulation or the presence of additional mechanisms that maintain the X chromosome in an inactive state, such as DNA methylation or histone deacetylation, in day 7 X-GFP-negative cells even after IFNγ-treatment (*23*, *25*, *54*). In summary, these data indicate that IFNγ treatment during reprogramming results in a higher activation of JAK-STAT3 signaling during early reprogramming, an increased expression of naive pluripotency genes and accelerated X-chromosome reactivation.

### Interferon γ treatment promotes active DNA demethylation in cells undergoing reprogramming

Global DNA demethylation is a hallmark of reprogramming to pluripotency in particular in female cells (*60*, *61*), and demethylation of X-chromosomal gene promoters is a critical step required for X-reactivation, although the demethylation mechanism of the X chromosome during reprogramming remains elusive (*23*). To gain further insight, we took advantage of mouse methylation BeadChip arrays (*62*) to study genome-wide and X-chromosomal 5-methylcytosine (5mC) and 5-hydroxy-methylcytosine (5hmC) levels, as 5mC is converted into 5hmC during active DNA demethylation (*63*). To assess the impact of IFNγ treatment (d0-5) on DNA demethylation during reprogramming, we analyzed the levels of 5mC and 5hmC in day 5 SSEA1+ and day 7 SSEA1+ X-GFP+ iPSCs, which is before and during the occurrence of X-reactivation, respectively (**Fig. 4D and 4E**).

We found that, in day 5 iPSC populations, IFNγ induced a general gain of the 5hmC mark on both autosomes and the X chromosome, globally and in all specific genomic regions analyzed (promoters, gene bodies and distal regions) (**Fig. 5A and 5B**), consistent with active DNA demethylation promoted by IFNγ. However, this did not result in detectable global differences in 5mC levels between control and IFNγ-treated iPSCs on day 5 (**Fig. S5A and S5B**). By contrast, in day 7 iPSCs, we observed a mild, but significant 5hmC increase specifically on X chromosomes but not in autosomes (globally, in promoters, gene bodies and distal regions) (**Fig. S5C and S5D**). Furthermore, we detected a global decrease of 5mC on day 7 IFNγ-treated iPSCs in all genomic regions analyzed (**Fig. 5C and 5D**). Importantly, the decrease in 5mC levels was stronger in X-chromosomal than in autosomal promoters. Together, these data suggest that IFNγ treatment early during reprogramming (d0-5) results in enhanced active DNA demethylation (increased 5hmC levels) on day 5, and a subsequent more efficient loss of 5mC at day 7 in X-reactivating iPSCs, with the 5mC loss being predominant in X-chromosomal promoters.

**Fig. 5.**
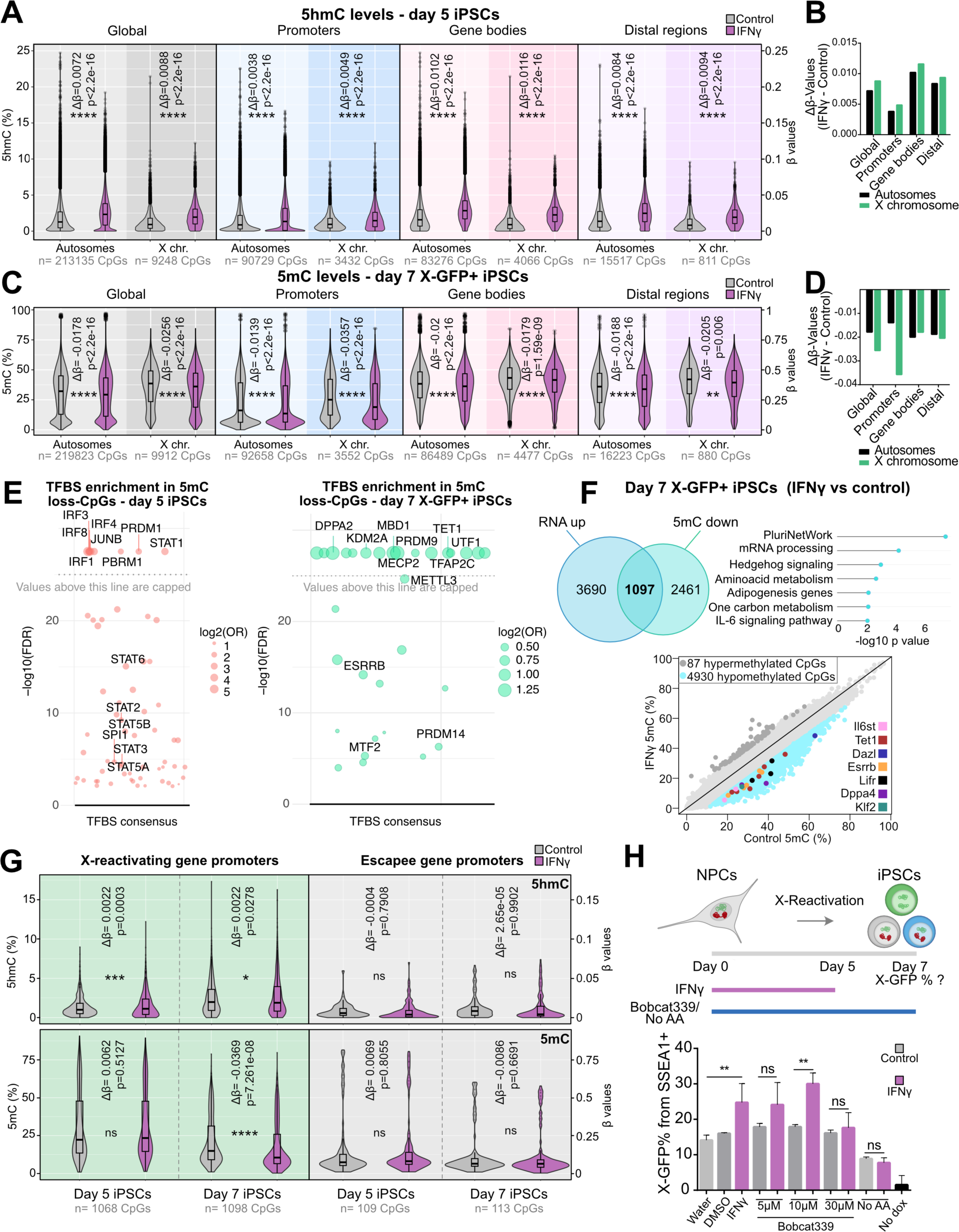
Interferon γ treatment promotes active DNA demethylation in cells undergoing reprogramming. (A) Analysis of 5hmC levels (β-values) of CpGs in autosomes and X chromosome in day 5 iPSCs for control and IFNγ conditions, globally and divided by genomic distribution: promoters (<= 1kb from TSS), gene bodies and distal regions (number (n) of detected CpGs from each category is indicated on the bottom of the graphs). Δβ-values (mean β-value IFNγ - mean β-value control) and p values (comparison IFNγ vs control) are shown in the graphs. Statistics: unpaired t-tests. (B) Δβ-values (mean β-value IFNγ - mean β-value control) for 5hmC in day 5 iPSCs for each genomic region (global, promoters, gene bodies and distal regions) in autosomes and X chromosome (corresponding to analysis in (A)). (**C**) Analysis of 5mC levels (β-values) of CpGs in autosomes and X chromosome in day 7 X-GFP+ iPSCs for control and IFNγ conditions, globally and divided by genomic distribution: promoters (<= 1kb from TSS), gene bodies and distal regions (number (n) of detected CpGs from each category is indicated on the bottom of the graphs). Δβ-values (mean β-value IFNγ - mean β-value control) and p values (comparison IFNγ vs control) are shown in the graphs. Statistics: unpaired t-tests. (**D**) Δβ-values (mean β-value IFNγ - mean β- value control) for 5mC in day 7 X-GFP+ iPSCs for each genomic region (global, promoters, gene bodies and distal regions) in autosomes and X chromosome (corresponding to analysis in (C)). (**E**) Transcription factor binding site (TFBS) enrichment analysis on differentially methylated CpGs (DMPs, logFC cutoff<(−0.1), p<0.01) which lose methylation upon IFNγ treatment compared to control in day 5 iPSCs (n=360 CpGs) and day 7 X-GFP+ iPSCs (n=10023 CpGs). Analysis was performed with Sesame R package. (**F**) Venn diagram (using Venny 2.1.0) representing overlapping of upregulated genes in the RNA-seq on the IFNγ X-GFP+ (medium or high) day 7 iPSCs (n=4787 upregulated genes) and genes associated with promoter DMPs (logFC cutoff <(−0.1), p<0.01) that lost 5mC upon IFNγ treatment (n=4930 CpGs corresponding to 3558 genes). Pathway enrichment of the 1097 common genes was analyzed with WikiPathways Mouse 2019. Scatter plot representing 5mC levels from promoter CpGs in day 7 IFNγ-treated and control iPSCs, showing hypermethylated CpGs (DMPs) in dark gray, hypomethylated CpGs (DMPs) in light blue, and selected pluripotency genes with significantly lower 5mC levels on their promoters. (**G**) Analysis of 5mC and 5hmC levels (β-values) of CpGs in X-reactivating (X-allelic ratio > 0.135, n=216 genes and 1068-1098 detected CpGs) and escapee (X-allelic ratio ≤ 0.135, n=20 genes and 109-113 detected CpGs) gene promoters at day 5 and day 7 X-GFP+ iPSCs for control and IFNγ conditions. Δβ-values (mean β-value IFNγ - mean β-value control) and p values (comparison IFNγ vs control) are shown in the graphs. Statistics: unpaired t-tests. (**H**) Experimental design: IFNγ treatment was performed from day 0-5 of reprogramming. Treatment with Bobcat339 (TET inhibitor) or absence of ascorbic acid (TET cofactor) was done during the 10 days of reprogramming. Flow cytometry analysis of X-GFP expression (from SSEA1+ cells) was performed on day 7 of reprogramming (n=3 technical replicates). IFNγ-TET inhibitor (Bobcat339, at 5 µM, 10 µM and 30 µM) and IFNγ-ascorbic acid absence during reprogramming were tested. Statistics (unpaired t tests): ns = non-significant; ** = p<0.01. Error bars represent SD.

Next, we analyzed transcription factor binding site (TFBS) enrichment in CpGs which showed a loss of 5mC on both days 5 and 7 of reprogramming upon early IFNγ treatment (**Fig. 5E**). On day 5 we observed an enrichment of binding sites corresponding to STAT1 (and other proteins from the STAT family such as STAT2 and STAT3) and IRF transcription factors, in line with the ongoing IFNγ treatment. In day 7 X-GFP+ iPSCs treated with IFNγ, we found an enrichment of binding sites corresponding mostly to transcription factors related to pluripotency, such as DPPA2, TFAP2C, UTF1, ESRRB and PRDM14, and epigenetic regulators (PRDM9, KDM2A, MBD1, TET1, MECP2, METTL3, MTF2). Most importantly, binding sites of TET1, the expression of which increased upon IFNγ treatment and catalyzes active DNA demethylation, showed up to be highly enriched in hypomethylated CpGs. In addition, we explored the overlap of the promoter CpGs which lost 5mC in IFNγ-treated day 7 X-GFP+ iPSCs with the genes gaining expression upon IFNγ treatment in the day 7 X-GFP+ populations (**Fig. 5F**). In total, up to 1097 common genes were found to lose 5mC in their promoter region and gain expression in this comparison. These genes were enriched in pathways such as the pluripotency network (*Tet1*, *Il6st*, *Dazl*, *Klf2*, *Esrrb*, *Lifr* and *Dppa4*), mRNA processing, metabolism and IL-6 signaling.

We then focused on the DNA methylation differences occurring on the X chromosome by analyzing the levels of 5hmC and 5mC in promoters of X-linked genes undergoing reactivation and in escapee genes, which are always active on the silenced X chromosome in differentiated cells (**Fig. 5G**). While we detected no changes for 5hmC or 5mC levels in escapee gene promoters, X-reactivating gene promoters showed a slight increase in 5hmC abundance in day 5 IFNγ-treated iPSCs, in line with the decreased levels of 5mC in day 7 X-GFP+ IFNγ-treated iPSCs. This increase in 5hmC and decrease in 5mC in X-reactivating gene promoters occurred specifically at X-linked genes which get reactivated later during reprogramming (“main”) (**Fig. S5F**) (*24*). Of note, 468 out of 470 differentially methylated X-chromosomal CpGs for 5mC showed a reduction in this mark (**Fig. S5G**), indicating that early IFNγ treatment (day 0-5) substantially boosts demethylation of the X-chromosome during its reactivation on day 7.

TET enzymes play a key role in active DNA demethylation (*63*), and are important for rewiring gene expression during pluripotency acquisition (*55*, *64–66*). Our gene expression analysis on days 5 and 7 of reprogramming showed that *Tet1* was overexpressed upon IFNγ treatment (**Fig. 3B,C,G**, **Fig. S3C,F**). As IFNγ treatment induces lower DNA methylation levels globally, and more pronouncedly on X-chromosomal promoters at day 7 of reprogramming, we wondered whether active DNA demethylation catalyzed by TET enzymes was responsible for the higher efficiency in X-chromosome reactivation upon IFNγ treatment. To functionally test this hypothesis, we induced reprogramming with or without ascorbic acid / vitamin C (cofactor enhancing TET enzyme activity and thereby iPSC reprogramming) (*67–69*) and with or without Bobcat339 (a TET inhibitor) (*70*), and we analyzed the levels of X-GFP reactivation by flow cytometry on day 7 of reprogramming (**Fig. 5H**). In the presence of ascorbic acid, IFNγ induced a higher percentage of X-GFP in comparison to the no IFNγ control condition (p=0.0026), consistent with our previous experiments (**Fig. 2C-E**). Without addition of IFNγ, the X-GFP percentage did not change upon Bobcat339 treatment, suggesting that in control conditions, TET enzymes might be dispensable for X-GFP-reactivation. While the addition of IFNγ together with low concentrations of Bobcat339 still induced a trend or significant increase in X-GFP percentage (5µM: p=0.16, 10µM: p=0.0026), this increase was no longer observed in the combination of IFNγ with the highest concentration of Bobcat339 (30 µM) (p=0.92), nor in the absence of ascorbic acid (p=0.27). This shows that, upon TET inhibition by Bobcat339 or by the absence of the TET-cofactor ascorbic acid, IFNγ treatment loses its ability to enhance X-chromosome reactivation. This suggests that the enhancing effect of IFNγ on X-reactivation is dependent on the catalytic activity of TET enzymes, indicating its potential mechanism of action.

## Discussion

In this study, we performed a genome-wide CRISPR knockout screen to identify genes and pathways involved in pluripotency and X-chromosome reactivation, which revealed both activators and repressors of these processes. We uncovered a role of the interferon γ pathway, the early activation of which during iPSC reprogramming results in a reduced colony number, while accelerating pluripotency acquisition and enhancing X-chromosome reactivation later on.

The decreased colony number induced by early IFNγ treatment could be caused by a reduced expression of focal adhesion genes and increased apoptosis during the first 2 days of reprogramming. In line with this, IFNγ treatment has been previously reported to disrupt β1 integrin-mediated focal adhesions in intestinal epithelial cells (*71*). Moreover, ADAM (a disintegrin and metalloproteinase) proteins have been found to act as reprogramming barriers by antagonizing focal adhesion through inhibition of specific integrin dimers (*43*), indicating an important role of focal adhesion during reprogramming. On the other hand, the accelerated pluripotency acquisition upon early IFNγ treatment during iPSC induction could be related to the observed increased STAT3 activation. IFNγ has been reported to induce activation of the STAT3 protein (and not only its canonical target STAT1) (*72*). STAT3, which is activated by the LIF signaling pathway, plays a key role in self-renewal of pluripotent stem cells (*73*) and induces the expression of pluripotency genes by binding to their regulatory elements together with OCT4, SOX2 and NANOG (*74*). Therefore, the enhanced activation of STAT3 induced by IFNγ could result in the higher expression of the pluripotency network earlier as observed from day 5 onwards, resulting in an acceleration of reprogramming. In line with this, a previous study demonstrated that constitutive activation of STAT3 induced a more efficient reprogramming, and inhibition of STAT3 signaling resulted in the absence of pluripotent colonies (*52*).

Another mediator of IFNγ pathway activation to accelerated reprogramming and/or X-reactivation could be the interferon regulatory factor 1 (IRF1). Overexpression of this transcription factor in porcine embryonic fibroblasts has been found to increase the efficiency of reprogramming to iPSCs through higher activation of the LIF-STAT3 pathway (*75*). In our study, IFNγ-induced *Irf1* expression peaked on day 2 of reprogramming, which could contribute to an increased expression of the pluripotency network, directly or through an enhanced STAT3 activation. Furthermore, the IFNγ-mediated enhancement of X-reactivation efficiency was disrupted in *Irf1* knockout cells. This shows that the increased and accelerated X-reactivation upon IFNγ pathway activation is, at least partially, dependent on IRF1.

The increased expression of the pluripotency network upon IFNγ treatment could also indirectly contribute to the observed enhanced X-chromosome reactivation. Pluripotency factors (e.g. OCT4, SOX2, NANOG and PRDM14) act as *Xist* repressors directly by binding to its intron 1 (*13*, *15*, *76*, *77*) and indirectly by repressing the *Xist*-activator *Rnf12/Rlim* (*14, 15*) and by activating the *Xist*-repressor *Tsix* (*76*, *78*). In our study, we observed a decreased *Xist* expression in IFNγ-treated cells on day 7 of reprogramming, which likely primed the cells for the enhanced X-reactivation. Even in day 7 X-GFP-negative cells, *Xist* expression was reduced after IFNγ treatment, but this was not sufficient to cause X-linked gene reactivation. This suggests the involvement of additional epigenetic silencing layers such as DNA methylation of X-chromosomal promoters to be present, which need to be removed for X-reactivation to take place (*23*, *25*, *54*, *79*, *80*).

DNA demethylation is a key step both for X-reactivation and for cellular reprogramming into iPSCs (*23*, *55*, *66*, *81–83*). Our findings revealed that IFNγ treatment induces the upregulation of *Tet1* and *Gadd45a*, which are known to play important roles in DNA demethylation (*84–86*). The expression of these genes increased from day 5 of reprogramming onwards, together with the upregulation of the pluripotency network. Our DNA (hydroxy)methylation analyses revealed that IFNγ treatment induced increased levels of 5hmC on day 5 of reprogramming, and decreased levels of 5mC at day 7 in cells undergoing X-reactivation. These results suggest that early treatment with IFNγ during reprogramming induces active DNA demethylation, which was preferentially happening at promoters corresponding to and/or bound by pluripotency factors, reflecting an acceleration in reprogramming upon IFNγ treatment. The loss of 5mC levels was more pronounced in X-chromosomal than in autosomal promoters specifically at X-linked genes undergoing reactivation, while this effect was not observed in escapee gene promoters, which are always active including on the silent X chromosome. A possible explanation for this X-specific effect of IFNγ treatment is that during female somatic cell reprogramming, DNA methylation erasure is more pronounced on the X chromosome than on autosomes due to higher DNA methylation levels on the inactive X chromosome.

Active DNA demethylation is mediated by the ten-eleven translocation (TET) enzymes (TET1, TET2 and TET3), which oxidize 5mC into 5-hydroxymethylcytosine (5hmC) (*86*). In this study we showed that in the presence of a TET inhibitor or in the absence of ascorbic acid (a TET cofactor that enhances TET activity (*68*, *87*) and is normally added to the medium in our reprogramming protocol), the IFNγ-mediated effect on X-reactivation disappeared, suggesting that IFNγ enhances X-reactivation in a TET-dependent manner. As IFNγ treatment induced an upregulation of *Tet1* expression from day 5 of reprogramming, higher levels of TET1 could be involved in the enhanced X-reactivation observed upon IFNγ pathway activation. Of note, in line with a previous study (*23*), the use of the TET inhibitor during reprogramming did not result in a lower efficiency of X-reactivation in the absence of IFNγ treatment. This indicates that TET-mediated active DNA demethylation is not needed for X-reactivation in a control reprogramming condition, although its enhancement by IFNγ treatment seems to boost the efficiency and kinetics of this process. By contrast, the absence of the TET-cofactor ascorbic acid during reprogramming decreased the efficiency of X-reactivation. This could be explained by the fact that ascorbic acid is not only a cofactor of TET enzymes, but also induces H3K9me2 and H3K36me2/3 demethylation by enhancing the activity of histone demethylases (*88*, *89*). As these histone marks are erased during iPSC reprogramming (*88*, *90*, *91*), this might be the reason why the absence of ascorbic acid during reprogramming, but not the addition of the TET inhibitor, had a detrimental effect in X-reactivation efficiency in the absence of IFNγ treatment.

Overall, our study revealed the IFNγ pathway as a novel player in iPSC reprogramming and X-chromosome reactivation, and that early activation of the pathway results in accelerated reprogramming and enhanced X-reactivation (**Fig. 6**). These findings provide new mechanistic insight into the process of X-reactivation and have potential impact on the reprogramming field, with the possibility to improve the generation of iPSCs. A recent study demonstrated that IFNγ promotes stemness in cancer cells (*92*), supporting the idea that the IFNγ pathway might be as well important for cellular dedifferentiation in other contexts highlighting the broader relevance of our findings. Although our study has been performed in the mouse model system, the X-chromosome status has been shown to be a sensitive measure of stem cell quality and differentiation potential of human female pluripotent cells (*93–96*). Therefore, a comprehensive understanding of the mechanisms regulating the X-chromosome state in both mouse and human and its link to pluripotency will be needed to improve the generation of stem cell lines suitable for disease modeling and clinical applications.

**Fig. 6.**
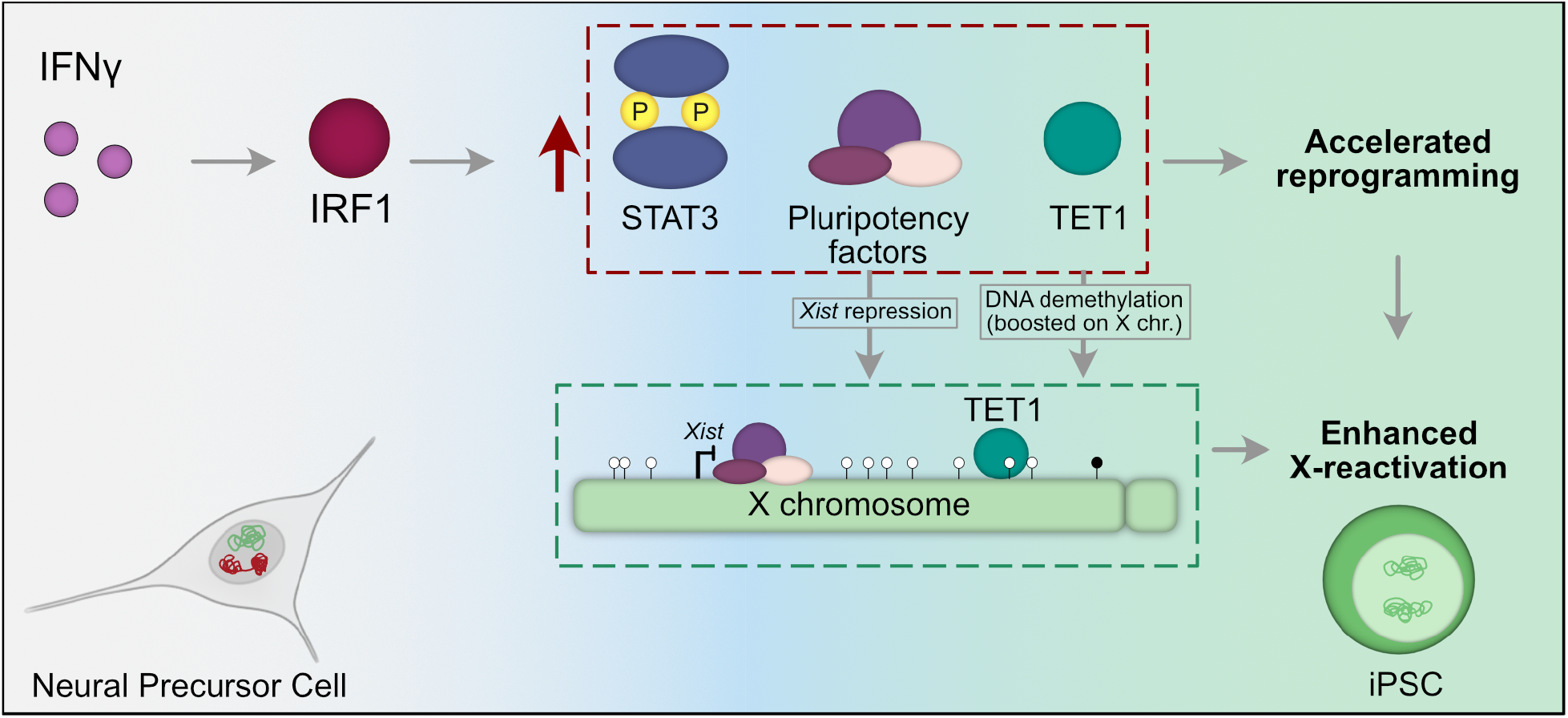
Model: Early activation of the IFNγ pathway impacts pluripotency acquisition and X-chromosome reactivation. The exposure to IFNγ in the early stages of NPC reprogramming into iPSCs induces the activation of IRF1 and a subsequent upregulation and activation of STAT3 and the expression of pluripotency genes, including *Tet1*. This would lead to an accelerated reprogramming kinetics. Moreover, the higher expression of pluripotency factors would lead to *Xist* repression, and higher levels of TET1 would induce a global loss of DNA methylation, which is most pronounced at X-chromosomal promoters of cells undergoing X-reactivation. This, together with the accelerated reprogramming, would explain the enhanced X-reactivation efficiency upon early IFNγ treatment during NPC reprogramming into iPSCs.

### Limitations of the study

X-chromosome reactivation is tightly linked to pluripotency. In fact, the expression of a robust pluripotency network is correlated with *Xist* repression (*13*, *97*). One important limitation of our CRISPR screen is that we did not identify genes or pathways playing a role exclusively in X-chromosome reactivation and not affecting pluripotency. Although the pluripotency reporter (*Nanog* promoter-RFP) from the PAX system (*24*) allows us to distinguish cells that only acquire late pluripotency from cells that also undergo X-reactivation, these populations were not included in the CRISPR screen due to the limited cell number showing these features and the high amount of cells needed for the screen. Therefore, we could identify pathways playing a role in both late pluripotency acquisition and X-reactivation, but we were not able to find genes or pathways that would uncouple these two processes. However, the fact that very few cells reached late pluripotency without undergoing X-reactivation indicates how closely related these two processes are.

## Materials and Methods

### Cell lines used

#### PAX cell line

As our starting cell line, we used the PAX (pluripotency and X-chromosome reporter) reprogramming system (*24*). The PAX system consists of a hybrid *Mus musculus* / *Mus castaneus* embryonic stem cell (ESC) line (*98*), in which the X-chromosome activity can be traced by the expression of an X-GFP reporter introduced into the *Hprt* locus of the *Musculus* X chromosome (X *mus*), which undergoes preferential inactivation when differentiated due to a truncation of the *Tsix* gene (*99*, *100*). This cell line also contains a Tet-On inducible *MKOS* (*cMyc* - *Klf4* - *Oct4* - *Sox2*) reprogramming cassette and a reverse tetracycline-controlled transactivator (rtTA) inserted into the *Sp3*-locus that allow iPSC induction from differentiated cells upon treatment with doxycycline (*49*). Moreover, it contains a pluripotency reporter (*Nanog* promoter-RFP or P-RFP) allowing the identification of cells that achieve a late pluripotent state during reprogramming.

#### PAX-iCas9 cell line

For the generation of the PAX-*iCas9* cell line, 5 million PAX ESCs were nucleofected with 3 μg of the Piggybac TRE-Cas9 plasmid, which was a gift from Mauro Calabrese (Addgene, Plasmid #126029) (*47*) and 3 μg of a Transposase plasmid kindly provided by Mitinori Saitou (*101*). The Amaxa Mouse Embryonic Stem Cell Nucleofector Kit was used (Lonza, VPH-1001), program A-24. Two days after transfection, cells were selected with 200 µg /ml of Hygromycin B Gold (Ibian tech., ant-hg-1) for 13 days, changing medium every day. Cells were single cell sorted by FACS using a BD FACSAria II and replated on 0.2% gelatin coated 96-well plates in serum-LIF medium with 200 µg/ml of Hygromycin B Gold. Colonies were expanded for 9 days and genotyped to detect the presence of the *Cas9* sequence. Genomic DNA was isolated from *iCas9*-transfected ESC clones (incubation at 55°C overnight with lysis buffer: 10% TrisHCl 1M pH 8, 5mM EDTA, 0.1% SDS, 0.2M NaCl in milliQ water). DNA was precipitated with isopropanol 1:1, and washed with EtOH 70%. The lysates were diluted 1:10. For PCR amplification, a DreamTaq PCR Master Mix was used (Thermo Fisher Scientific, K1082). To functionally test the knockout production efficiency of selected clones, a gRNA targeting *GFP* was cloned into a Lenti-guide puro plasmid (a gift from Feng Zhang (Addgene plasmid #52963; http://n2t.net/addgene:52963; RRID:Addgene_52963)) (*102*). 293T cells were transfected with the plasmids pCMVR8.74 (a gift from Didier Trono (Addgene plasmid #22036; http://n2t.net/addgene:22036; RRID:Addgene_22036), pCMV-VSV-G (a gift from Bob Weinberg (Addgene plasmid #8454; http://n2t.net/addgene:8454; RRID:Addgene_8454))(*103*)and the Lenti guide puro-*GFP* gRNA plasmid. Viral harvesting and concentration was performed 48 hours post transfection using the Lenti X Concentrator (Clontech, 631231), following the manufacturer’s instructions. The PAX-*iCas9* ESC clones were infected with lentiviruses containing the *GFP* gRNA, and the virus was removed after 24 hours. 48 hours post-infection, ESC media containing 2 µg/ml of puromycin (Ibian Tech., ant-pr-1) was added to the cells. Cells were exposed to puromycin for 4 days, prior to treatment with doxycycline for 6 days and measuring the percentage of X-GFP+ cells by flow cytometry every day, using a BD LSRFortessa Cell Analyzer.

#### Stat1−/− and Irf1−/− cell lines

gRNA pairs targeting the *Stat1* or *Irf1* gene, or a scramble gRNA, were cloned into a Lenti-guide puro (a gift from Feng Zhang (Addgene plasmids #52963; http://n2t.net/addgene:52963; RRID:Addgene_52963)) or Lenti-guide blast (a gift from Brett Stringer (Addgene plasmid #104993; http://n2t.net/addgene:104993; RRID:Addgene_104993)) plasmids (*102*, *104*). 293T cells were thawed and maintained in DMEM medium (Thermo Fisher Scientific, 31966021) for 5 days. The day before transfection, 20 million 293T cells were seeded on one 150 mm plate per gRNA. The next day, 293T cells were transfected with 7.5 μg of of the plasmid pCMVR8.74 (a gift from Didier Trono (Addgene plasmid #22036; http://n2t.net/addgene:22036; RRID:Addgene_22036), 3 µg of the plasmid pCMV-VSV-G (a gift from Bob Weinberg (Addgene plasmid #8454; http://n2t.net/addgene:8454; RRID:Addgene_8454)) (*103*) and 10 µg of the Lenti guide puro/blast-gRNA plasmid using 1 mg/ml of PEI transfection reagent (Tocris, 7854). Incubation with the transfection mix was done for 5 hours at 37 °C, and media was replaced for 25 ml of viral harvest media per 150 mm plate (DMEM medium with 30% FBS and 100U/ml penicillin/streptomycin). Viral harvesting was performed 48 hours post transfection, followed by filtering with 0.45µm PES filters. Viruses were concentrated by using the Lenti X Concentrator (Clontech, 631231), following the manufacturer’s instructions. The PAX-*iCas9* ESC line was infected with lentiviruses containing the gRNA pairs, and virus was removed after 24 hours. 48 hours post-infection, ESC media containing 2 µg/ml of puromycin (Ibian Tech., ant-pr-1) and/or 5 µg/ml of blasticidin (Ibian Tech., ant-bl-1) was added to the cells. Cells were exposed to puromycin for 4 days and to blasticidin for 6 days. Then, cells were treated with doxycycline for 7 days, followed by single cell sorting, PCR screening of clones to detect the presence of gRNA pairs and western blot to detect the absence of IRF1 or STAT1 protein.

#### Feeders (irradiated Mouse Embryonic Fibroblasts)

Mouse Embryonic Fibroblasts were obtained from E12.5 mouse embryos and expanded for 10 days at 37 °C with 5% CO2 and 5% O2 in DMEM medium (Thermo Fisher Scientific, 31966021) supplemented with 10% FBS (Thermo Fisher Scientific, 10270106), 25mM HEPES (Thermo Fisher Scientific, 15630056), 1mM Sodium Pyruvate (Thermo Fisher Scientific, 11360070), 1x MEM NEAA (Thermo Fisher Scientific, 11140050), 50U/ml penicillin/streptomycin (Ibian Tech, P06-07100) and 0.1mM 2-mercaptoethanol (Thermo Fisher Scientific, 31350010), prior to gamma irradiation (30 kGy) for inactivation. Mouse care and procedures were conducted according to the protocols approved by the Ethics Committee on Animal Research of the Parc de Recerca Biomedica de Barcelona (PRBB) and by the Departament de Territori i Sostenibilitat of the Generalitat de Catalunya (Ref. No. 10469).

### Embryonic stem cell culture

Mouse embryonic stem cells were cultured at 37 °C with 5% CO2 on 0.2% gelatin-coated plates in serum/LIF medium: DMEM medium (Thermo Fisher Scientific, 31966021) supplemented with 10% FBS (ES pre-tested, Capricorn, FBS-ES-12A), 1,000 U/ml LIF (ORF Genetics, 01-A1140-0100), 25mM HEPES (Thermo Fisher Scientific, 15630056), 1mM Sodium Pyruvate (Thermo Fisher Scientific, 11360070), 1x MEM NEAA (Thermo Fisher Scientific, 11140050), 50U/ml penicillin/streptomycin (Ibian Tech, P06-07100) and 0.1mM 2-mercaptoethanol (Thermo Fisher Scientific, 31350010). Medium was changed every day. Passaging of cells was done using 0.05%

Trypsin-EDTA (Thermo Fisher Scientific, 25300054). PCR mycoplasma tests were performed monthly.

### Neural precursor cell differentiation

Neural precursor cell (NPC) differentiation and reprogramming were done similar as in Bauer et al (*24*). Mouse ESCs were thawed on serum/LIF medium 5 days before induction, and passaged for 3 consecutive days onto 0.2% gelatin coated plates at 1,75 × 10^5^ cells per cm^2^. The day of induction, media was changed to 2i/LIF: 50% Neurobasal medium (Thermo Fisher Scientific, 12348017), 50% DMEM F12 (Thermo Fisher Scientific, 21041025), 1x N2 (Thermo Fisher Scientific, 17502048), 1x B27 (Thermo Fisher Scientific, 12587001), 3 µM CHIR99021 (Sigma Aldrich, SML1046), 0.4 µM PD0325901 (Selleck Chemicals, S1036) and LIF 1,000 U/ml (ORF Genetics, 01-A1140-0100). After 6 hours, cells were dissociated with Accutase (Merck Millipore, SF006) and plated on 0.2% gelatin coated T75 flasks at a density of 6,666 cells/cm^2^ in RHBA medium (Takara Bio, Y40001). Media was changed every 2 days. From day 6, media was supplemented with 10 ng/ml EGF (R&D Systems, 236-EG-200) and 10 ng/ml bFGF (Thermo Fisher Scientific, 13256029). From day 8 onwards, media was also supplemented with ROCK inhibitor 10 µM (Selleck Chemicals, S1049). On day 9 of differentiation, cells were dissociated with Accutase (Merck Millipore, SF006) and incubated with anti-SSEA1 micro beads (Miltenyi Biotec, 130-094-530) at 4°C for 15 minutes. MACS separation was performed in order to enrich for SSEA1 negative cells. Staining with SSEA1 eFluor 660 antibody 1:50 (Thermo Fisher Scientific, 50-8813-42) was performed at 4°C for 15 minutes. A BD FACSAria II SORP was used to sort the SSEA1 negative, P-RFP negative, X-GFP negative cells. 1.5 × 10^6^ sorted cells were plated on a 0.2% gelatin coated well of 6-well plate in RHBA supplemented with EGF, bFGF and ROCK inhibitor. Media was changed every day until day 12.

### Reprogramming of neural precursor cells into induced pluripotent stem cells

At day 12 of NPC differentiation, the NPC differentiation media (RHBA with EGF, bFGF and ROCKi) was supplemented with 25 mg/ml L-ascorbic acid (Sigma Aldrich, A7506) and 1 mg/ml doxycycline (Tocris, 4090/50). One day later, cells were dissociated with Accutase (Merck Millipore, SF006) and seeded at different densities depending on day of analysis (49,100 cells per cm^2^ for day 5, 12,300 cells per cm^2^ for day 7 and 2,850 cells per cm^2^ for day 10) on top of male irradiated Mouse Embryonic Fibroblasts (iMEFs) on 0.2% gelatin coated plates in iPSC medium: DMEM medium (Thermo Fisher Scientific, 31966021), 15% FBS (ES pre-tested, Capricorn, FBS-ES-12A), 25mM HEPES (Thermo Fisher Scientific, 15630056), 1mM Sodium Pyruvate (Thermo Fisher Scientific, 11360070), 1x MEM NEAA (Thermo Fisher Scientific, 11140050), 50U/ml penicillin/streptomycin (Ibian Tech, P06-07100) and 0.1mM 2-mercaptoethanol (Thermo Fisher Scientific, 31350010), supplemented with 1,000 U/ml LIF, 25 mg/ml L-ascorbic acid and 1 mg/ml doxycycline. Media was changed on days 3, 5, 7, 8 and 9.

### Lentiviral CRISPR KO screen

#### gRNA library amplification

The gRNA library used for the screening was the Mouse Improved Genome-wide Knockout CRISPR Library v2 (a gift from Kosuke Yusa, Addgene, #67988) (*48*), with 90,230 gRNAs targeting 18,424 genes (average of 5 gRNAs per gene). NEB 10-beta Electrocompetent *E. coli* (NEB, C3020K) were electroporated in five concomitant reactions (each reaction containing 20 µL of bacteria and 1 µL of the gRNA library (20 ng/µl). After electroporation, 1 ml of SOC recovery medium was added to each reaction and bacteria were incubated at 37 °C for 1 hour shaking. Bacteria were then grown overnight at 37 °C shaking in 1L 2×TY (5 g/l NaCl, 16 g/l tryptone, 10 g/l yeast extract) + ampicillin 100 µg/ml. The plasmid gRNA library was purified by using the QIAfilter Plasmid Maxi Kit (Qiagen, 12263), following the manufacturer’s instructions. Concentration was measured with Nanodrop (Thermo Fisher Scientific, ND-1000).

#### Generation of lentiviral gRNA library

For the lentiviral library production, 293T cells were thawed in DMEM medium (Thermo Fisher Scientific, 31966021) supplemented with 10% FBS (Thermo Fisher Scientific, 10270106). After 2 and 4 days, cells were passaged into 3 and 5 T175 flasks, respectively (2.5 × 10^6^ cells and 40 ml of media per flask). At day 7, cells were seeded on 10 T175 flasks for transfection at a density of 18 million cells and 25 ml of media per T175 flask. After 24 hours, transfection was done by using 31 µg of the plasmid library, 38.8 µg of the plasmid pCMVR8.74 (a gift from Didier Trono (Addgene plasmid #22036; http://n2t.net/addgene:22036; RRID:Addgene_22036), 3.88 µg of the plasmid pCMV-VSV-G (a gift from Bob Weinberg (Addgene plasmid #8454; http://n2t.net/addgene:8454; RRID:Addgene_8454)) (*103*), 6 ml of OPTIMEM (Thermo Fisher Scientific, 11058021) and 305 µL of TransIT-LT1 Transfection Reagent (Mirus Bio, MIR 2300) per T175 flask. Incubation with the transfection mix was done for 8 hours at 37 °C, and media was replaced for 60 ml of viral harvest media per T175 flask (DMEM medium with 30% FBS and 100U/ml penicillin/streptomycin). Viral harvesting was performed 36 hours post transfection, followed by filtering with 0.45µm PES filters. Viruses were concentrated by using the Lenti X Concentrator (Clontech, 631231), following the manufacturer’s instructions.

#### gRNA library Lentiviral infection of ESCs

*iCas9*-PAX ESCs were thawed in ESC medium and amplified for 3 days. 13 T175 flasks coated with 0.2% gelatin were seeded with 18.5 × 10^6^ ESCs per flask, in 27 ml of ESC medium with 8 µg/ml Polybrene (Merck, TR-1003-G) and the lentiviral gRNA library. The next day, media was replaced with ESC medium containing 2 µg/ml of puromycin (Ibian Tech., ant-pr-1). Media with antibiotics was replaced every other day for 1 week. In parallel, 72 hours post-infection, the percentage of BFP positive cells was measured using a BD LSR Fortessa flow cytometer, in order to calculate the MOI (0.06) and the coverage (200 cells per gRNA). gRNA sequencing was performed to check gRNA representation.

#### NPC differentiation, reprogramming and cell isolation by FACS

For the CRISPR screening, two independent biological replicates (each one with two technical replicates) were performed in different differentiation and reprogramming inductions. To this end, 1,2 × 10^8^ pooled lentiviral-infected ESCs were thawed on 3 150 mm plates in serum/LIF medium 5 days before induction, and passaged 3 days in a row onto 0.2% gelatin coated plates at a density of 25 million cells per 150 mm plate (4 plates). The day of induction, media was changed to 2i/LIF for 6 hours, and cells were then dissociated with Accutase (Merck Millipore, SF006) and seeded on 52 gelatin-coated T75 flasks at a density of 7,5 × 10^5^ cells per flask in RHBA medium. Differentiation was followed as previously described. Sorting of SSEA1-P-RFP-X-GFP-NPCs was performed on day 9, as described above. 4 × 10^7^ NPCs were sorted in the Replicate 1, and 8,4 × 10^7^ NPCs were sorted in the Replicate 2. Each 1.5 × 10^6^ sorted cells were plated on a 0.2% gelatin coated well of 6-well plate in RHBA supplemented with EGF, bFGF and ROCK inhibitor. Media was changed every day until day 12. Cell pellets of 2×10^6^ cells were collected for gRNA abundance analysis.

For reprogramming, 6,6 × 10^8^ mouse male feeders were thawed on 60 gelatin-coated 150 mm plates one day prior to reprogramming induction, in DMEM medium (Thermo Fisher Scientific, 31966021) supplemented with 10% FBS (Thermo Fisher Scientific, 10270106), 25mM HEPES (Thermo Fisher Scientific, 15630056), 1mM Sodium Pyruvate (Thermo Fisher Scientific, 11360070), 1x MEM NEAA (Thermo Fisher Scientific, 11140050), 50U/ml penicillin/streptomycin (Ibian Tech, P06-07100) and 0.1mM 2-mercaptoethanol (Thermo Fisher Scientific, 31350010). At day 12 of neural precursor cell differentiation, the NPC differentiation media (RHBA with EGF, bFGF and ROCKi) was supplemented with 25 mg/ml L-ascorbic acid and 1 mg/ml doxycycline. One day later, cells were dissociated with Accutase (Merck Millipore, SF006) and seeded on 46 (Replicate 1) and 60 (Replicate 2) 150 mm plates on top of male irradiated Mouse Embryonic Fibroblasts (iMEFs) (3,000 cells per cm^2^) in iPSC medium supplemented with 1,000 U/ml LIF, 25 mg/ml L-ascorbic acid and 1 mg/ml doxycycline. Media was changed on days 3, 5, 7, 8 and 9. At day 10 of reprogramming, cells were dissociated with 0.25% trypsin EDTA (Thermo Fisher Scientific, 25200056). Trypsinization was stopped with DMEM 10% FBS containing 10 µg/ml of DNAse I (Sigma Aldrich, 11284932001). Cells were then stained with SSEA1 eFluor 660 antibody 1:100 (Thermo Fisher Scientific, 50-8813-42) at 4 °C for 45 minutes in rotation. A BD FACSAria II SORP was used to sort three different populations, according to the BFP fluorescence (gRNA plasmid), SSEA1-APC fluorescence (pluripotency marker) and X-GFP (X Chromosome status): Non-Pluripotent population (BFP+ SSEA1-X-GFP-), Early Pluripotent population (BFP+ SSEA1+ X-GFP-) and Late Pluripotent, X-Chromosome Reactivated population (BFP+ SSEA1+ X-GFP+). Cell pellets were collected and frozen at −80 °C until processed for gDNA extraction.

#### Sample preparation and guide-DNA sequencing

Genomic DNA was extracted from cell pellets using the DNeasy Blood and Tissue Kit (Qiagen, 69504). 1.15 × 10^7^NPCs and 1.4 × 10^7^ cells of each reprogramming population were processed for Replicate 1, and 2.64 × 10^7^ NPCs and 1.8 × 10^7^ cells of each reprogramming population were processed for Replicate 2. For amplification of the gRNAs and introduction of the Illumina-sequencing adapters, two consecutive PCRs were performed by using the Q5 High-Fidelity DNA Polymerase (NEB, M0491). For the PCR 1, all the extracted gDNA was used for amplification, in PCR reactions of 50 µL with 1 µg of gDNA as template. For this PCR 1, all forward primers and all reverse primers were mixed together in the “Forward primer mix” and “Reverse primer mix” in equal amounts to have a final concentration of 10 uM (1.67 uM of each primer). *Sequences follow the IUPAC nucleotide code.* After electrophoresis in a 2% agarose gel, the DNA was purified by using a QIAEX II Gel Extraction Kit (Qiagen, 20051). For PCR 2, 16 reactions of 50 µL were performed per sample, by using 5 ng of the purified PCR 1 product as template for each reaction. Independent PCR reactions for each sample were done with reverse primers containing different barcodes for sample identification. Electrophoresis in a 2% agarose gel was performed prior to DNA purification from gel. PCR components and quantities are indicated in **Table 1**. PCR conditions are specified in **Table 2**. Sequencing was performed using an Illumina HiSeq 2500 (50 bp single-end).

**Table 1.** PCR components for gRNA library amplification.

| Component | Volume for 50 µl reaction |
| --- | --- |
| **PCR1** |  |
| 5x Q5 buffer | 10 µl |
| dNTP 10 mM | 1 µl |
| F mix 10 µM (Stag0_F to Stag5_F, 1.67 µM each) | 2.5 µl |
| R mix 10 µM (Stag0_R to Stag5_R, 1.67 µM each) | 2.5 µl |
| H2O | To 50 µl |
| DMSO | 3 µl |
| Q5 polymerase | 0.5 µl |
| DNA | 1 µg |
| **PCR2** |  |
| 5x Q5 buffer | 10 µl |
| dNTP 10 mM | 1 µl |
| F primer 10 µM (TS-HT-D5x-1-F) | 2.5 µl |
| R primer 10 µM (different for each sample) | 2.5 µl |
| H2O | To 50 µl |
| Q5 polymerase | 0.5 µl |
| DNA (PCR1 product) | 5 ng |

**Table 2.** PCR conditions for gRNA library amplification.

| Temperature | Time | Cycles |
| --- | --- | --- |
| **PCR1** |  |  |
| 98°C | 3 min | - |
| 98°C | 30 seconds | X 20 |
| 56.5°C | 20 seconds |  |
| 72°C | 60 seconds |  |
| 72°C | 2 min | - |
| 4 °C | Hold | - |
| **PCR2** |  |  |
| 98°C | 3 min | - |
| 98°C | 30 seconds | X 8 |
| 56.5°C | 20 seconds |  |
| 72°C | 60 seconds |  |
| 72°C | 2 min | - |
| 4 °C | Hold | - |

**Table 3.**
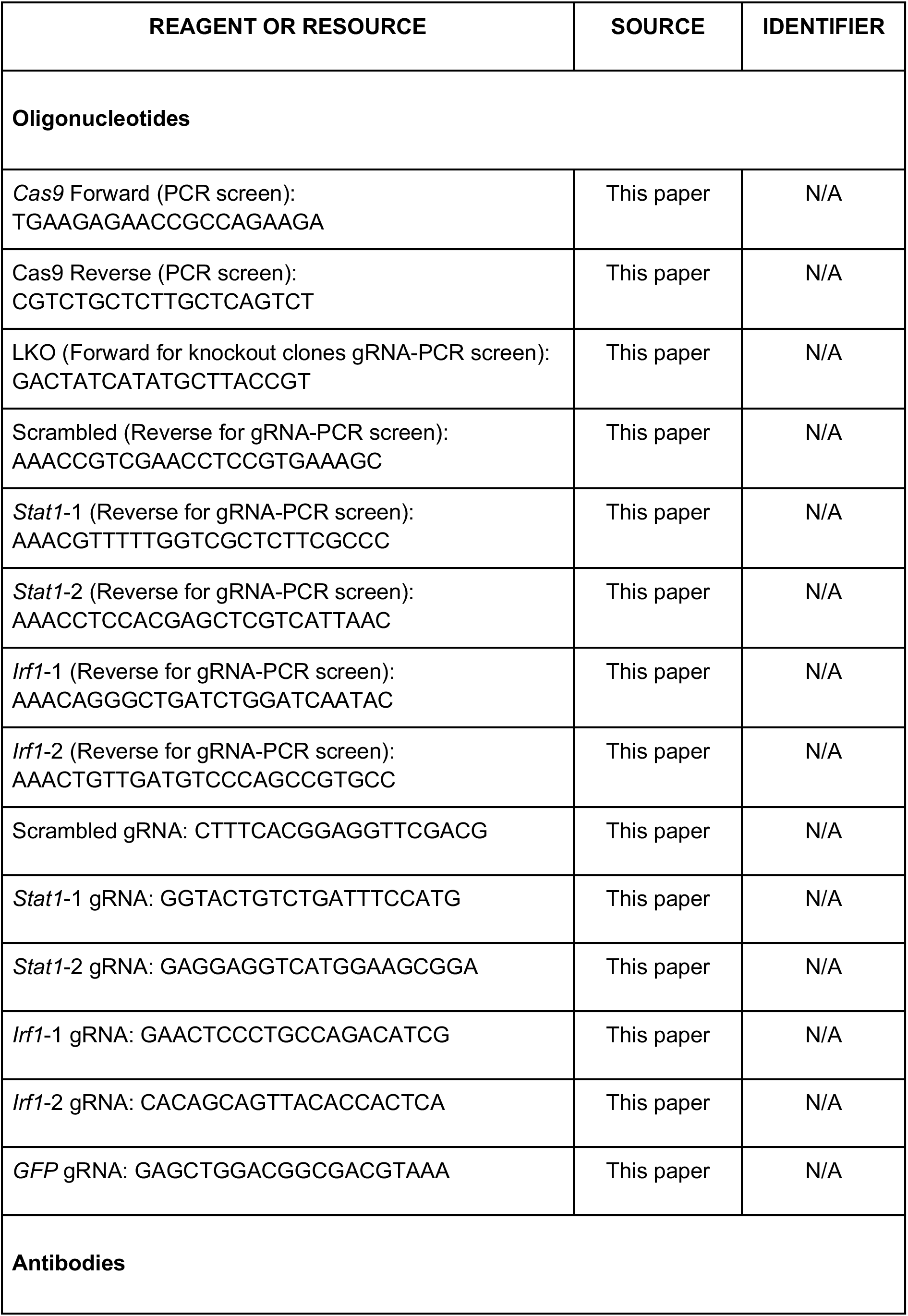
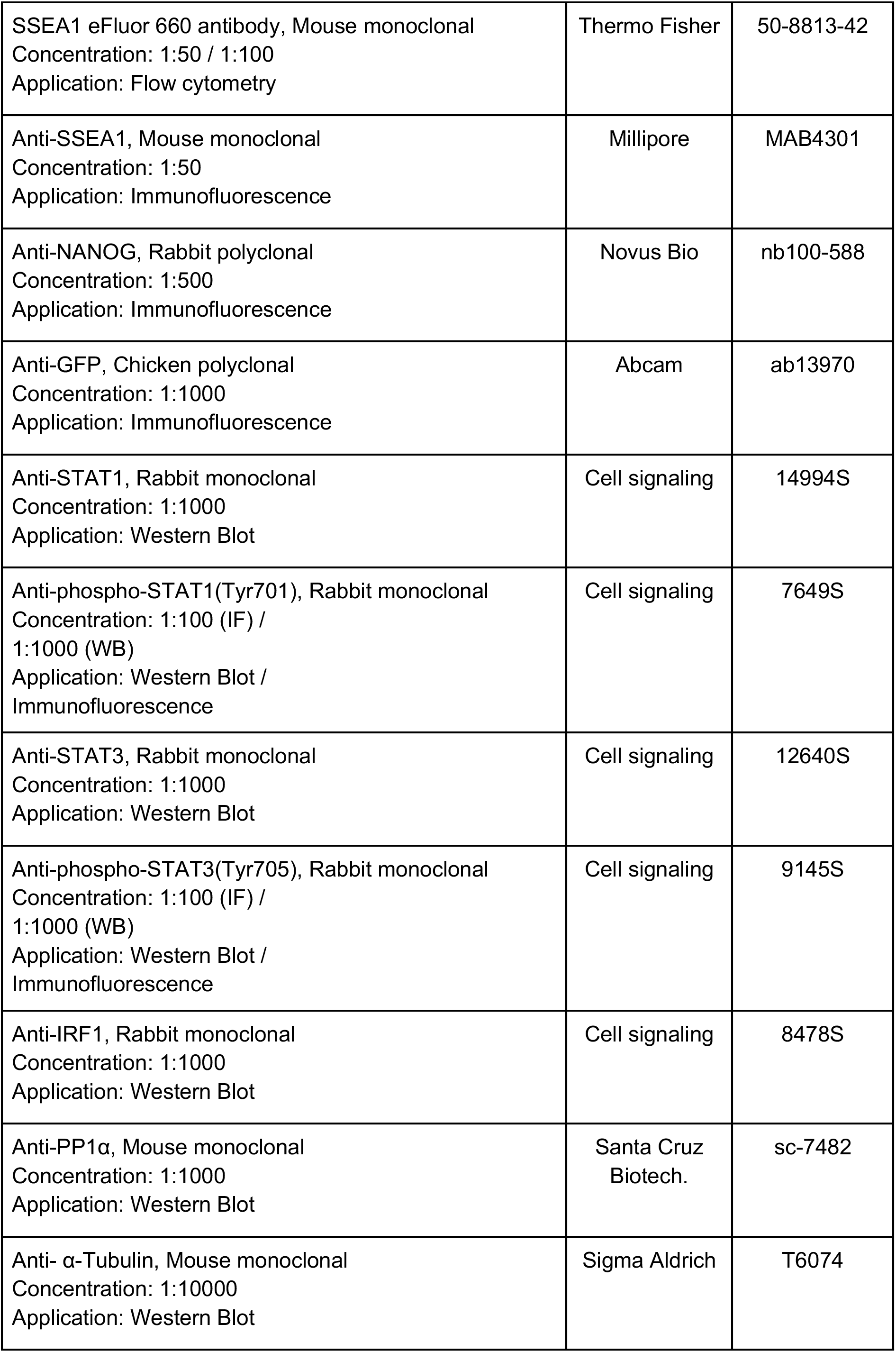
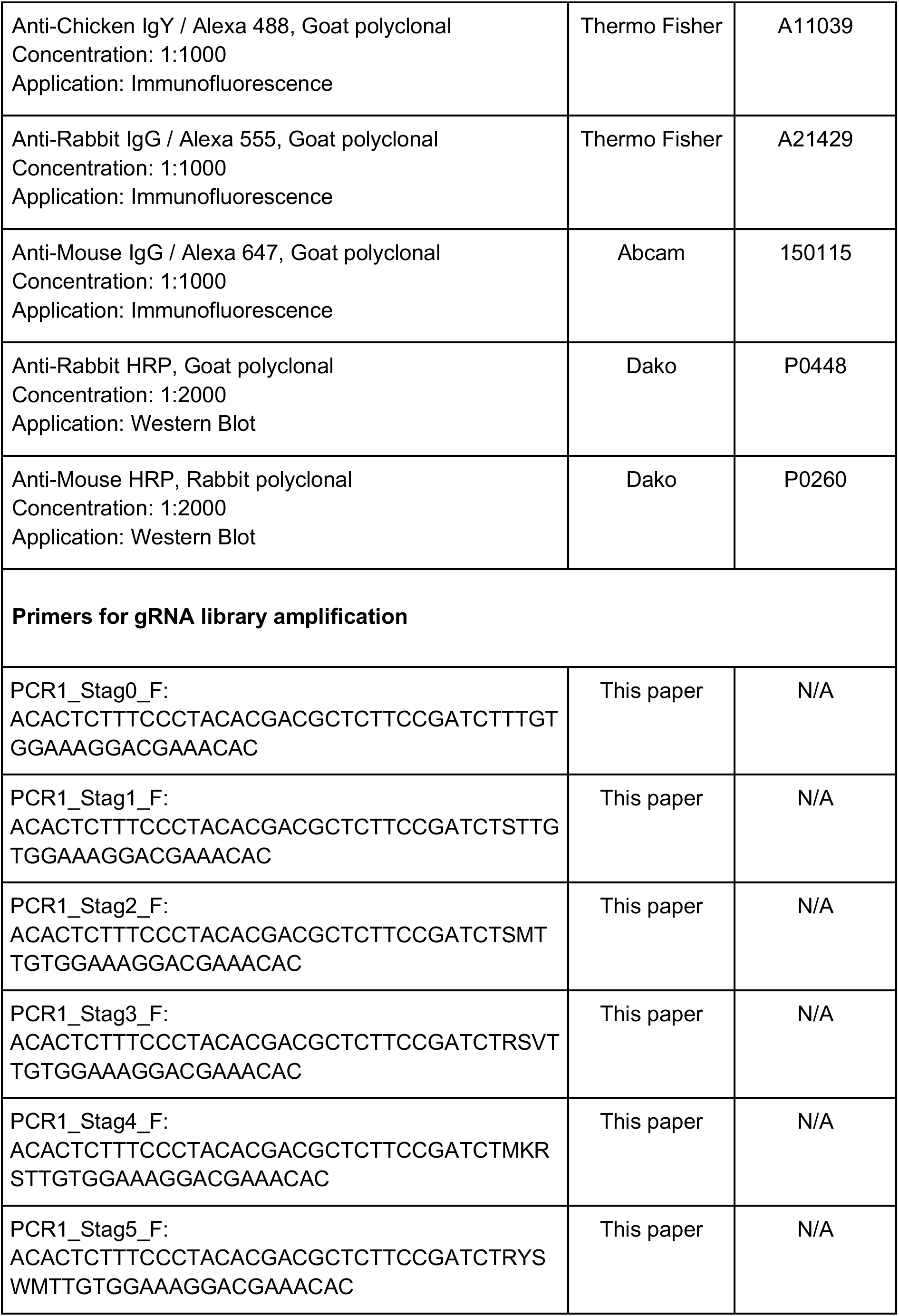
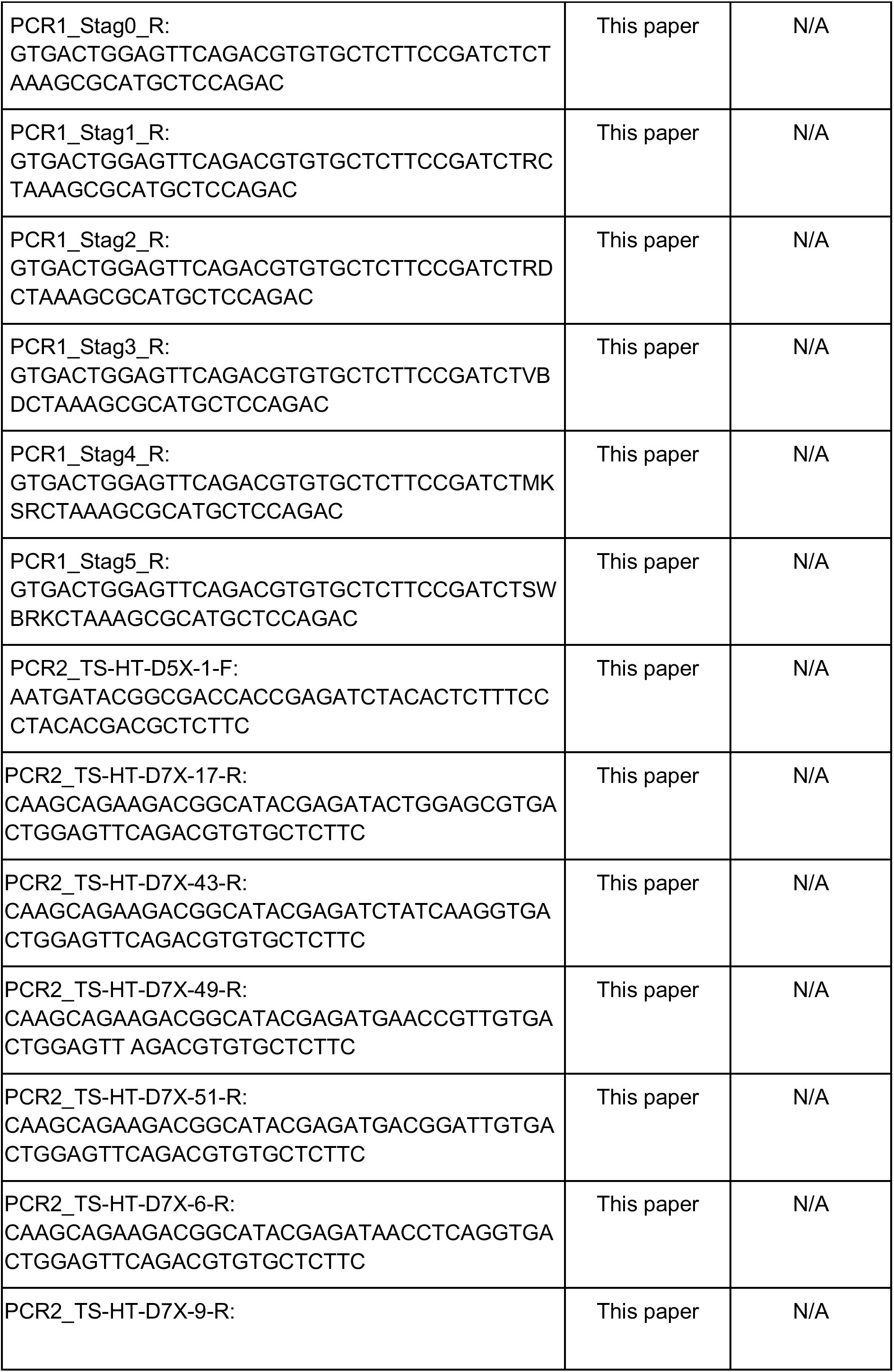
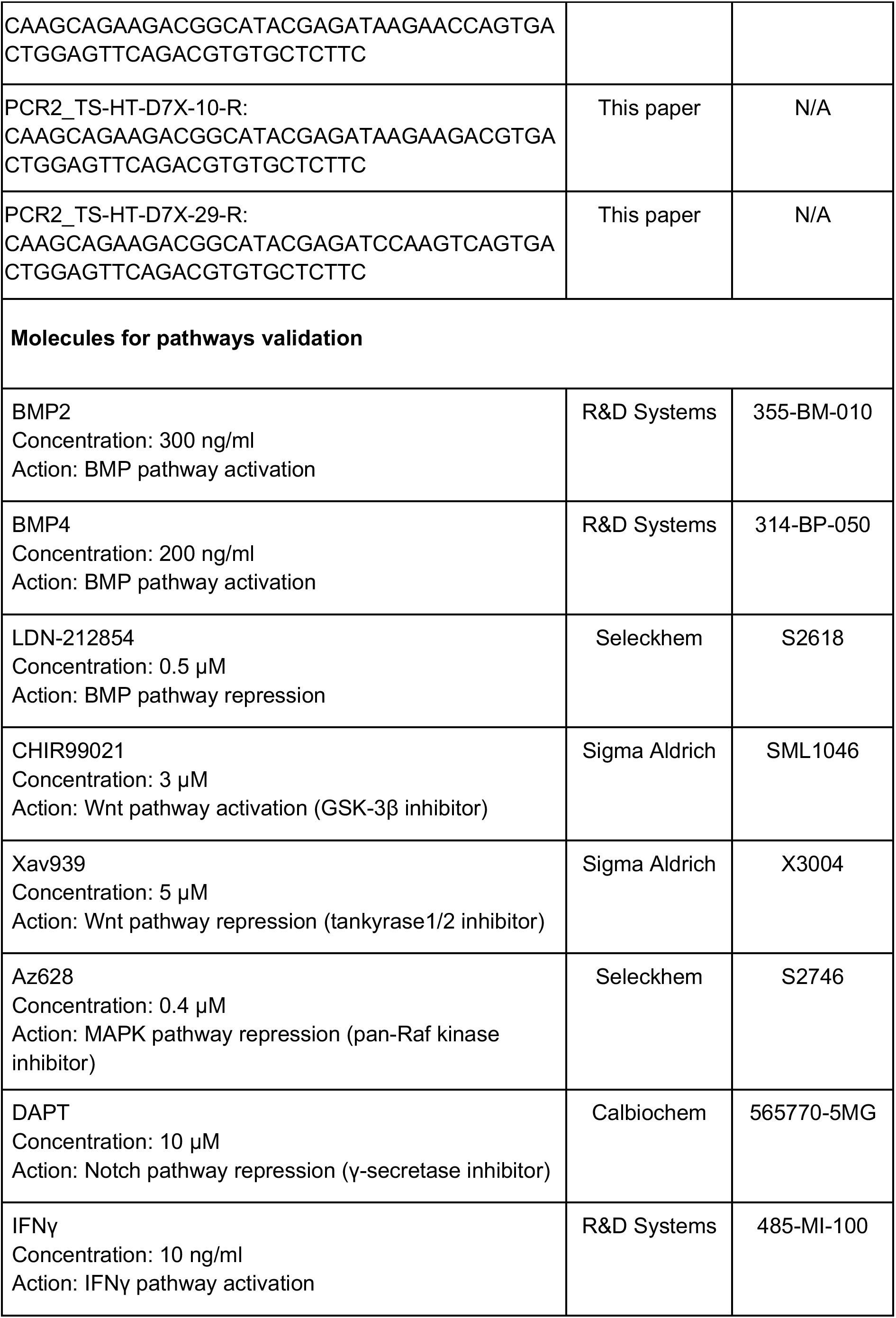
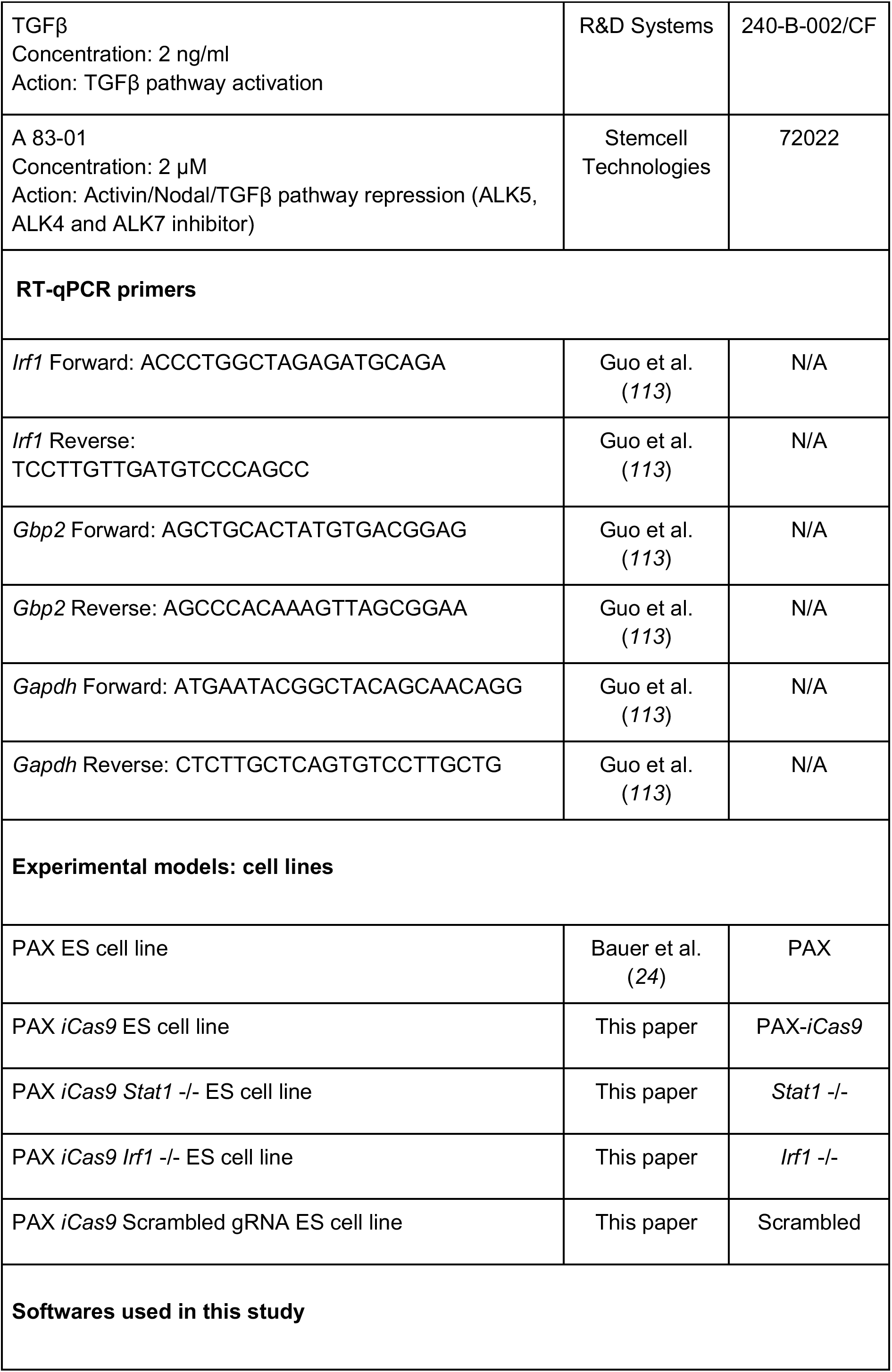
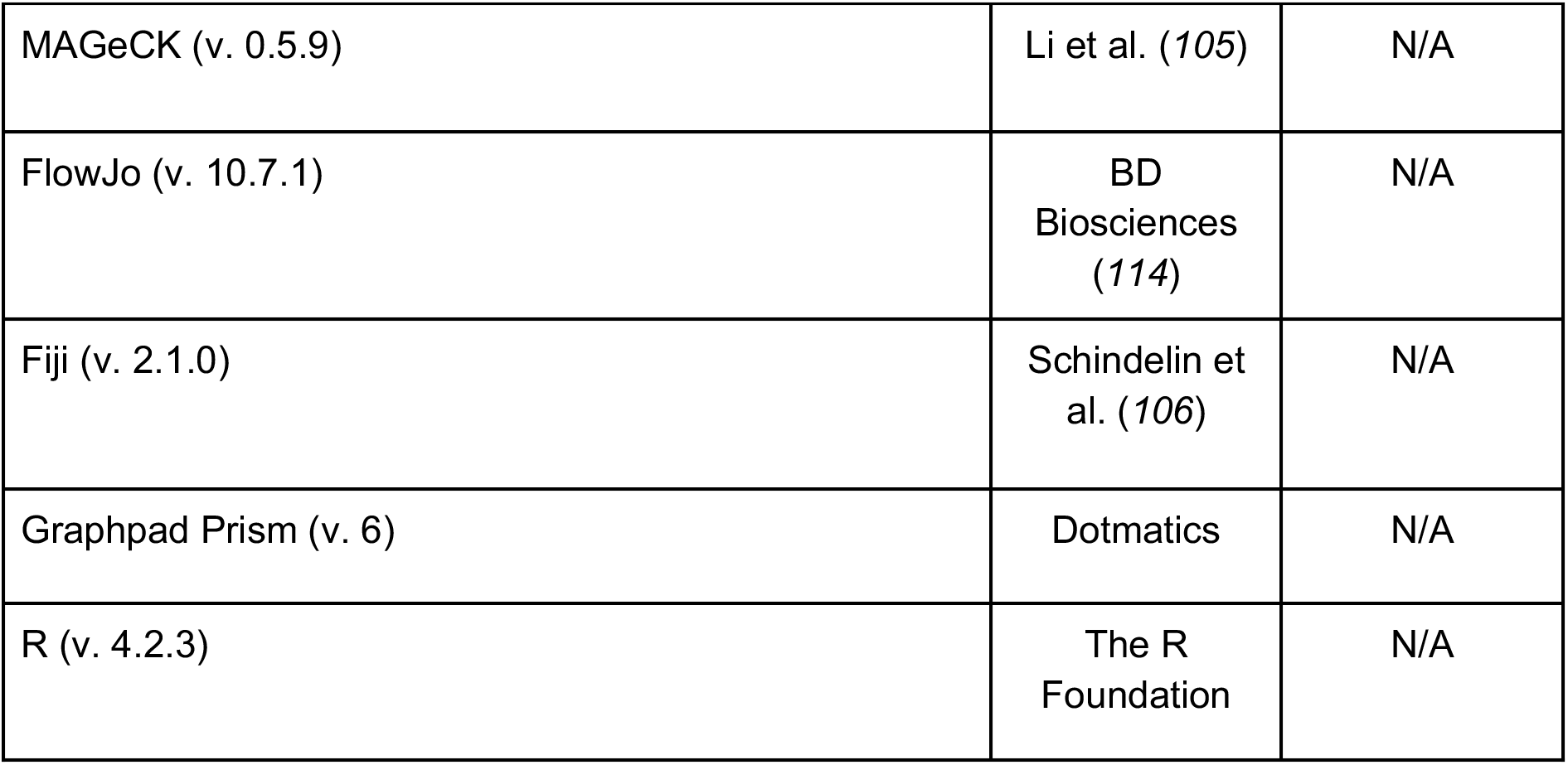
Resources Table.

### CRISPR screening analysis

The gRNA sequencing from the CRISPR KO screening was analyzed with MAGeCK software (*105*). The gRNA abundance of each population was determined by taking into account the two biological replicates (with two technical replicates each). The gRNA abundance comparisons were performed pairwise. The list of overrepresented genes for the comparison of the reprogramming populations (non-pluripotent, early pluripotent and late pluripotent) to NPCs was obtained by selecting the top 250 genes of each comparison ranked by positive score in the MAGeCK software and filtering for unique genes from the obtained list. The list of essential genes (underrepresented for the comparison of each reprogramming population to NPCs) was obtained by filtering common genes with an RRA score <0.05 and L2FC of <(−0.75). For the pairwise comparisons among the reprogramming populations (early pluripotent vs non-pluripotent, late pluripotent vs early pluripotent), the selection of hits was performed by using an RRA score of <0.05, a L2FC of <(− 0.8) and a “goodsgrna” equal or higher than 3. Gene Ontology pathways enrichment analysis was performed with the obtained filtered genes using the library “WikiPathways Mouse 2019” in the Enrichr website (https://maayanlab.cloud/Enrichr/).

### CRISPR screening pathways validation – molecule screening

For the molecular screening, NPC differentiation was performed as previously described. At day 13 of NPC differentiation, cells were dissociated with accutase (Merck Millipore, SF006) and seeded on top of male irradiated Mouse Embryonic Fibroblasts in iPSC medium supplemented with 1,000 U/ml LIF, 25 mg/ml L-ascorbic acid and 1 mg/ml doxycycline. Three seeding densities were used: 49,100 cells per cm^2^ for analysis at day 5 of reprogramming, 12,300 cells per cm^2^ for analysis at day 7, and 2,850 cells per cm^2^ for analysis at day 10. Media was changed on days 3, 5, 7 and 9. Molecules were added from day 0 to 5, from day 5 to 10, and from day 0 to 10, for all the conditions. At days 5, 7 and 10, cells were dissociated with 0.25% trypsin EDTA (Thermo Fisher Scientific, 25200056) and stained with SSEA1 eFluor 660 antibody 1:100 (Thermo Fisher Scientific, 50-8813-42) at 4 °C for 30 minutes in rotation. A BD LSRFortessa Cell Analyzer was used to check the SSEA1-APC fluorescence (early pluripotency marker) and X-GFP (X-chromosome status).

### Flow cytometry

A BD FACSAria II SORP was used for cell sorting. Neural precursor cells were sorted at around 3,500 events per second, maximum flow rate of 4 with a 100 µm nozzle was used to increase cell viability after sorting. SSEA1 negative P-RFP negative X-GFP negative cells were selected. iPSCs were sorted at around 8,000 events per second, using the 85 µm nozzle, selecting the cell populations regarding SSEA1-APC and X-GFP fluorescence. A BD LSRFortessa Cell Analyzer or a BD LSR II Flow Cytometer were used for Flow Cytometry Analysis experiments. For CRISPR screening experiments, FVS780 (BD Horizon, 565388) was used as a viability dye at 1.1 ng/mL. For the rest of the experiments, DAPI (Biotium, BT-40043) was used at 0.1 μg/ml.

### Interferon γ treatment during iPSC reprogramming

Recombinant Mouse IFNγ Protein (R&D Systems, 485-MI-100) was added to the iPSC medium at a concentration of 10 ng/ml from day 0-5, 5-10 or 0-10. Further analysis was performed by flow cytometry using a BD LSRFortessa Cell Analyzer, alkaline phosphatase staining, immunofluorescence or cell sorting by a BD FACSAria II SORP for western blotting, RNA-sequencing or DNA methylation arrays.

### Interferon γ treatment during NPC differentiation

Recombinant Mouse IFNγ Protein (R&D Systems, 485-MI-100) was added to the NPC medium at a concentration of 10 ng/ml from day 0-5, 5-10 or 0-10. At days 5 and 10, cells were dissociated with accutase and stained with SSEA1 eFluor 660 antibody 1:100 (Thermo Fisher Scientific, 50-8813-42) at 4 °C for 30 minutes on ice. A BD LSRFortessa Cell Analyzer was used to check the SSEA1-APC fluorescence (early pluripotency marker) and X-GFP (X-chromosome status).

### RNA isolation, cDNA synthesis and quantitative RT-PCR

Total RNA isolation was performed with the RNeasy Plus Mini Kit (Qiagen, 74136) or RNeasy Micro Kit (Qiagen, 74004). Concentration was quantified with Nanodrop (Thermo Fisher Scientific, ND-1000). cDNA was synthesized using a High-Capacity RNA-to-cDNA Kit (Thermo Fisher Scientific, 4387406). qRT-PCR was performed in triplicates for each sample, using Power SYBR Green PCR Master Mix (Thermo Fisher Scientific, 4367659). Gene expression levels were calculated as 2^(−ΔCT) normalized with the average CT of the housekeeping gene *Gapdh*.

### DNA methylation modifiers experiments

Reprogramming was done as described previously, combining the IFNγ (10 ng/ml, day 0 to 5) (R&D Systems, 485-MI-100) with the addition of the TET inhibitor molecule Bobcat 339 (concentration of 5 µM, 10 µM or 30 µM, R&D Systems, 6977/10) or in the absence of ascorbic acid (normally added to the reprogramming medium at 25 mg/ml, Sigma Aldrich, A7506) from day 0 to 7. 4,3 × 10^4^ NPCs were plated per well of a 12-well plate. On day 7 of reprogramming, cells were detached from the plates using 0.25% Trypsin-EDTA (Thermo Fisher Scientific, 25300054) and stained with SSEA1-APC antibody (1:100) and DAPI. Analysis was done using a BD LSRFortessa Cell Analyzer.

### Apoptosis assay

Day 13 NPCs (treated with doxycycline and ascorbic acid during 24 hours) were induced for reprogramming on CFSE-stained feeders (to sort these cells out; 0.5 µM of CFSE CellTrace, Thermo Fisher Scientific, C34554), stained and plated the day before on gelatin-coated plates, 1 × 10^6^ feeders seeded per well of a 6-well plate). 2 × 10^5^ NPCs were seeded per well of a 6-well plate, in iPSC medium in the absence or presence of IFNγ (R&D Systems, 485-MI-100) at 10 ng/ml. Three experimental replicates were done. After 48h, cells were detached from the plates using 0.05% Trypsin-EDTA (Thermo Fisher Scientific, 25300054) and stained with Annexin V-APC antibody and DAPI (0.1 μg/ml, Biotium, BT-40043) using the Annexin V Apoptosis Detection Kit (ThermoFisher, 88-8007-72). Analysis was done using a BD LSR II Flow Cytometer.

### Western blotting

For protein extraction, cells were resuspended in Laemmli Buffer and boiled at 95°C for 10 minutes. Protein extracts were loaded in a 10% acrylamide gel (BioRad, 1610149) and electrophoresis was performed for protein separation. Transference was done into a PVDF membrane (Sigma Aldrich, P2938). Blocking of the membrane was performed using 4% milk in TBS 0,5% Tween 20 (Sigma Aldrich, P7949) for 1 hour at room temperature. The membrane was incubated overnight at 4°C with the corresponding antibodies (rabbit anti STAT1 1:1000 (Cell Signaling, 14994S), rabbit anti phospho STAT1 Tyr701 1:1000 (Cell Signaling, 7649S), mouse anti PP1α 1:1000 (Santa Cruz Biotech., sc-7482), rabbit anti STAT3 1:1000 (Cell Signaling, 12640S), rabbit anti phospho STAT3 Tyr705 1:1000 (Cell signaling, 9145S), rabbit anti IRF1 1:1000 (Cell signaling, 8478S), mouse anti α-Tubulin 1:10000 (Sigma Aldrich, T6074) in blocking solution. Secondary antibody incubation was performed in polyclonal rabbit anti mouse – HRP antibody 1:2000 (Dako, P0260) or polyclonal goat anti rabbit – HRP antibody 1:2000 (Dako, P0448) in blocking solution for 1.5h at room temperature. For washes, TBS 0.5% Tween 20 was used. The membranes were developed by using an EZ-ECL kit (Reactiva, 120500120) and X-ray films (Rosex Medical, EWPJH).

Fiji (*106*) was used to calculate the relative intensities of STAT3 and pSTAT3 Tyr705. The “mean gray value” of each Regions Of Interest (ROI) with the same area was calculated for both loading controls and proteins of interest, followed by the inversion of the pixel density and the calculation of the ratio for each sample.

### Alkaline phosphatase staining

Cells were washed with PBS and fixed with 4% PFA for 10 minutes at room temperature. Washing with milliQ water was done before adding the Alkaline Phosphatase staining solution (10 ml of milliQ water, 10 mg of Fast red TR salt hemi (zinc chloride) salt (Chem Cruz, sc-215025) and 400 µL of Naphthol AS-MX phosphate (Sigma-Aldrich, 855-20ML)) for 10 minutes. Cells were washed again with water and medium was aspirated prior to scanning.

### Immunofluorescence

Cells were seeded on 0.2% gelatin-coated coverslips on 12-well plates, where the reprogramming experiments were performed. On the specific reprogramming day, coverslips were washed with PBS and fixed at RT for 10 minutes with 4% PFA (Electron Microscopy Sciences, 15713S). After fixation, samples were washed with PBS and permeabilized with PBS 0.5% Triton (Sigma Aldrich, T8787) for 10 minutes at RT. Then, coverslips were washed once with 70% EtOH and kept in 70% EtOH at −20°C until staining was performed. For immunostaining, coverslips were washed in PBS and blocking was performed with PBS 2% BSA (Sigma Aldrich, SLCK2178) 0.2% triton for 1.5 hours at RT. Primary antibody incubation was done overnight at 4°C (rabbit anti pSTAT1 Tyr701 (Cell signaling, 7649S), rabbit anti pSTAT3 Tyr705 (Cell signaling, 9145S), mouse anti SSEA1 (Sigma Aldrich, MAB4301), rabbit anti NANOG (Novus Bio, nb100-588), chicken anti GFP (Abcam, ab13970)). Coverslips were washed three times with PBS for 5 minutes at RT, and secondary antibody incubation was performed for 2 hours at RT (goat anti mouse A647 (Abcam, 150115), goat anti rabbit A555 (Thermo Fisher, A21429), goat anti chicken A488 (Thermo Fisher, A11039)). Coverslips were washed three times with PBS for 5 minutes at RT, adding DAPI 10 µg/ml (Biotium, BT-40043) in the last wash. Mounting was done with Vectashield Antifade Mounting Medium (Vector Laboratories, H-1000-10). Images were taken with a Leica SP5 confocal/MP inverted microscope.

### RNA-sequencing experiments

For the RNA-seq of day 2 iPSCs, male irradiated Mouse Embryonic Fibroblasts were stained with

0.5 μM CFSE CellTrace (Thermo Fisher Scientific, C34554) and seeded on 0.2% gelatin-coated plates upon thawing. One day after, day 13 NPCs were seeded on top of the CFSE-stained feeders in iPSC medium to induce reprogramming, in presence or absence of IFN γ (R&D Systems, 485-MI-100, 10 ng/ml). On day 2 of reprogramming, CFSE negative cells were sorted by using a BD FACSAria II SORP, and cell pellets were kept at −80 °C until RNA extraction.

For the RNA-seq of iPSCs on days 5 and 7, day 13 NPCs were seeded on top of feeders in iPSC medium to induce reprogramming, in presence or absence of IFN γ (R&D Systems, 485-MI-100, 10 ng/ml) for the first 5 days. On days 5 and 7, cells were dissociated with 0.25% trypsin EDTA (Thermo Fisher Scientific, 25200056). Trypsinization was stopped with DMEM 10% FBS containing 10 µg/ml of DNAse I (Sigma Aldrich, 11284932001). Cells were then incubated with anti-SSEA1 micro beads (Miltenyi Biotec, 130-094-530) at 4 °C for 15 minutes. MACS separation was performed in order to enrich for SSEA1 positive cells. Staining with SSEA1 eFluor 660 antibody 1:100 (Thermo Fisher Scientific, 50-8813-42) was performed at 4 °C for 45 minutes. Sorting was performed by using a BD FACSAria II SORP. For iPSCs at day 5, SSEA1 positive cells were sorted. For iPSCs at day 7, SSEA1 positive cells were separated into three populations: X-GFP negative, medium and high cells. Cell pellets were kept at −80 °C until RNA extraction.

RNA was extracted from cell pellets by using a RNeasy Plus Mini Kit (Qiagen, 74136) or RNeasy Micro Kit (Qiagen, 74004). Concentration was quantified with Nanodrop (Thermo Fisher Scientific, ND-1000).

### RNA-seq analysis

The RNA libraries preparation was performed by ribosomal RNA depletion using the TruSeq Stranded Total RNA Library Preparation Kit (Illumina, 20020596). Sequencing was performed by an Illumina HiSeq 2500 (50 bp paired-end or 125 bp paired-end, merged and trimmed to 50 bp for further analysis). RNA sequencing analysis was done similar as in Severino et al (*11*). FastQ files passing the quality control were aligned to the mm10 reference genome, which contained CAST/EiJ and 129S1/SvImJ SNPs positions masked. The SNPs positions of the mouse strains were obtained from ftp://ftp-mouse.sanger.ac.uk/REL-1505-SNPs_Indels/mgp.v5.merged.snps_all.dbSNP142.vcf.gz.tbi. A VCF file containing only the SNPs positions from CAST/EiJ and 129S1/SvImJ strains was generated. Alignment of reads to the reference genome was done using STAR (*107*) with implementation of the WASP method (van de Geijn et al. 2015) to filter allele specific alignments. The output BAM files were used to obtain the read counts using the HTseq tool (v0.6.1) (*108*). These steps were performed using a published Nextflow pipeline (*109*) and following the workflow described in https://github.com/biocorecrg/allele_specific_RNAseq. Around 75-85% of reads aligned to the reference genome, corresponding to 3,5 × 10^7^ −5 × 10^7^ mapped reads. Differential expression analysis was done with the R package DESeq2 (v1.32.0) (Love, Huber, and Anders 2014). Differentially expressed genes were identified performing pairwise comparisons. Read counts were normalized by library size and filtered for having a mean across the samples >10. Log2 fold change shrinking was done with the “normal” parameter. Upregulated and downregulated genes were selected by filtering for a positive or negative Log2FC (respectively) and an adjusted p value of <0.1 (for control vs IFNγ comparisons in all timepoints). Gene Ontology pathways enrichment analysis was performed with the obtained filtered genes using the library “WikiPathways Mouse 2019” in the Enrichr website (https://maayanlab.cloud/Enrichr/).

To run the principal component analysis, we employed the top 500 genes showing highest variability. ggplot2 R package (v3.3.5, https://ggplot2.tidyverse.org) was used for generating the heatmap (representing the Z score of FPKM of selected genes in the different cell populations). To calculate the pluripotency score, the expression levels of *Nanog*, *Zfp42*, *Dppa4*, *Dppa5a*, *Esrrb*, *Prdm14* and *Sall4* in each time point were normalized to the expression of these genes in the ESCs, and the average of these values was represented for two independent replicates.

For the allelic ratio analysis, 315 protein-coding genes that showed over 25% of total X-linked gene expression in the X cas in all the populations were selected. To calculate the X mus proportion, we divided the X mus expression to the sum of X mus and X cas expression (X mus / (X mus + X cas)) in the selected genes.

### DNA (hydroxy)methylation experiments and analyses

Reprogramming was induced in the presence or absence of IFNγ at 10 ng/ml (from day 0 to 5) (R&D Systems, 485-MI-100). SSEA1+ day 5 iPSCs (4 replicates from different reprogramming rounds) and SSEA1+ X-GFP+ day 7 iPSCs (2 replicates from different reprogramming rounds) were sorted with a BD FACSAria II SORP. DNA was extracted using Wizard® Genomic DNA Purification Kit (Promega, A1120). After measuring DNA quantity by Qubit (ThermoFisher Scientific), 2µg of each sample was evenly splitted for the oxidation reaction (Oxidative Bisulfite - OxBS-treated samples) and the mock-oxidation reaction (Bisulfite-BS-treated samples) where the oxidant solution was replaced by water following the TrueMethyl oxBS Module manufacturer’s instructions (NuGEN-Tecan, 0414). Both aliquots were then processed in parallel for all stages of the protocol. After the oxidation reaction where 5-hydroxymethylcytosine is oxidized to 5-formylcytosine (5fC) and 5-methylcytosine (5mC) stays unchanged, the bisulfite treatment converts 5fC and all non-methylated cytosines to uracil, while 5mC is not altered.

For samples to be run on the Illumina Infinium® Mouse Methylation BeadChip Array (Illumina, 20041558), 7 μL of recovered TrueMethyl template were mixed with 1 μL of 0.4 N NaOH following manufacturer’s instructions. All subsequent steps were completed following the Infinium HD Assay Methylation protocol (https://emea.support.illumina.com/content/dam/illumina-support/documents/documentation/chemistry_documentation/infinium_assays/infinium_hd_methylation/infinium-hd-methylation-guide-15019519-01.pdf).

The DNA methylation status of the studied samples was obtained using the Infinium Mouse Methylation BeadChip Array (∼285,000 methylation sites). GenomeStudio Software 2011 (Illumina) was used to process raw signal intensities. The mm10 mouse genome manifest from Illumina was used as reference, as described in the Illumina manifest file associated with the Infinium Mouse Methylation BeadChip. DNA methylation β values were obtained from raw IDAT files using the software’s default normalization with control probes and background subtraction. The 5mC signal was extracted from the β values of the OxBS samples, while the 5hmC signal was obtained by subtracting the β values of the BS samples from those of the OxBS samples. All further analyses were performed using the R environment (v4.2.3). To remove erratic probe signals, quality control steps were applied. Probes that did not pass the intensity threshold were removed (intensity < 1000), as well as those with detection p value > 0.01. 5mC and 5hmC levels were then batch-corrected using the Limma R package (v3.50.3) (*110*).

Differentially (hydroxy)methylated positions ((h)DMPs) were extracted using the function topTable from the limma package (v3.50.3), adjusting by Benjamini-Hochberg method. CpGs with p values < 0.01 were selected, and further filtering with log fold change (logFC) was also performed (logFC +/− 0.1). The package ggplot2 (v3.3.5, https://ggplot2.tidyverse.org) was used to create the heatmap of the DMPs. In order to assign the genomic features corresponding to each CpG, ChIPseeker package (v1.30.3) (*111*) together with org.Mm.eg.db (v3.14.0) for annotation were used. The distribution violin and box plots were generated with ggplot2. Gene Ontology pathways enrichment analysis was performed with the obtained filtered genes using the library “WikiPathways Mouse 2019” in the Enrichr website (https://maayanlab.cloud/Enrichr/). The overlap analysis of DMPs and RNA-seq differentially expressed genes was done using Venny 2.1.0 (https://bioinfogp.cnb.csic.es/tools/venny/).

For selection of X-linked reactivating or escapee genes, protein-coding genes that showed over 25% of total X-linked gene expression in the X cas in all the populations analyzed from the RNA-seq dataset and showed an allelic ratio under 0.135 in NPCs (X-reactivating genes) or above 0.135 in NPCs (escapee genes) were selected (similarly as in (*11*, *24*)). The lists of “early” and “main” X-reactivating genes were obtained from (*24*).

Transcription factor binding site (TFBS) enrichment was analyzed with the Sesame R package (v1.16.1) (*112*).

### Statistical analyses

For experiments with technical replicates, unpaired t-tests were performed. For experiments with independent reprogramming rounds, paired t-tests were done. In the molecule validation experiments, paired t-tests were performed. A confidence interval of 95% was used. Each molecule was compared to their diluent control: BMP2, BMP4, TGFβ and IFNγ were compared to water, while the rest of the molecules were compared to DMSO controls. Each treatment was compared to its specific time point (day 0-5, day 5-10, day 0-10). For allelic ratio and DNA methylation data comparisons, unpaired t-tests were performed.

## Acknowledgments

We thank present and previous Payer laboratory members for discussions and feedback. We are grateful to C. Arnan for sharing CRISPR screen expertise; T. Graf and S. Sdelci for advice on the CRISPR screen follow-up; M. Maqueda for assistance on the DNA methylation analysis with Sesame; L. Cozzuto for help with the allele-specific pipeline and T. Mattimoe and T. Graf for critical reading of the manuscript. We also acknowledge critical technical support by the CRG core facilities including the CRG Genomics Unit; the CRG/UPF FACS Unit; the Advanced Light Microscopy Unit; the Bioinformatics Unit and the PRBB Animal Facility. We would like to thank the Spanish Ministry of Economy, Industry and Competitiveness (MEIC), to the EMBL partnership and to the “Centro de Excelencia Severo Ochoa”. We also acknowledge support of the CERCA Programme of the Generalitat de Catalunya.

## Funding

Agencia Estatal de Investigación (AEI) of the Spanish Ministry of Science and Innovation grant BFU2017-88407-P (BP)

Agencia Estatal de Investigación (AEI) of the Spanish Ministry of Science and Innovation grant EUR2019-103817 (BP)

Agencia Estatal de Investigación (AEI) of the Spanish Ministry of Science and Innovation grant PID2021-123383NB-I00 (BP)

Agencia Estatal de Investigación (AEI) of the Spanish Ministry of Science and Innovation grant PID2019-111243RA-I00 (JLS)

AXA Research Fund (BP)

Agencia de Gestió d’Ajuts Universitaris i de Recerca (AGAUR), 2017 grant SGR 346 (BP) Agencia de Gestió d’Ajuts Universitaris i de Recerca (AGAUR) 2021 grant SGR 01222 (BP) Instituto de Salud Carlos III grant CP19/00176 (JLS)

Ramón Areces Foundation (Doctoral Thesis in Life Sciences Fellowship) (M. Barrero) La Caixa International PhD Fellowship Program (M. Bauer)

## Author contributions

Conceptualization: M. Barrero, M. Bauer, BP Methodology: M. Barrero, AL, ALR, EB, LGP Investigation: M. Barrero, AL, ALR, EB, LGP Visualization: M. Barrero

Supervision: BP, JLS, LDC, AB Writing—original draft: M. Barrero, BP

Writing—review & editing: M. Barrero, BP, JLS, M. Bauer, EB

## Competing interests

Authors declare that they have no competing interests.

## Data availability

Raw and processed data generated will be available upon peer-reviewed publication.

## Supplemental Information

The Interferon γ Pathway Enhances Pluripotency and X-Chromosome Reactivation in iPSC Reprogramming

Mercedes Barrero, Anna V. López-Rubio, Aleksey Lazarenkov, Enrique Blanco, Moritz Bauer, Luis G. Palma, Anna Bigas, Luciano Di Croce, José Luis Sardina and Bernhard Payer.

---

**Figure S1.**
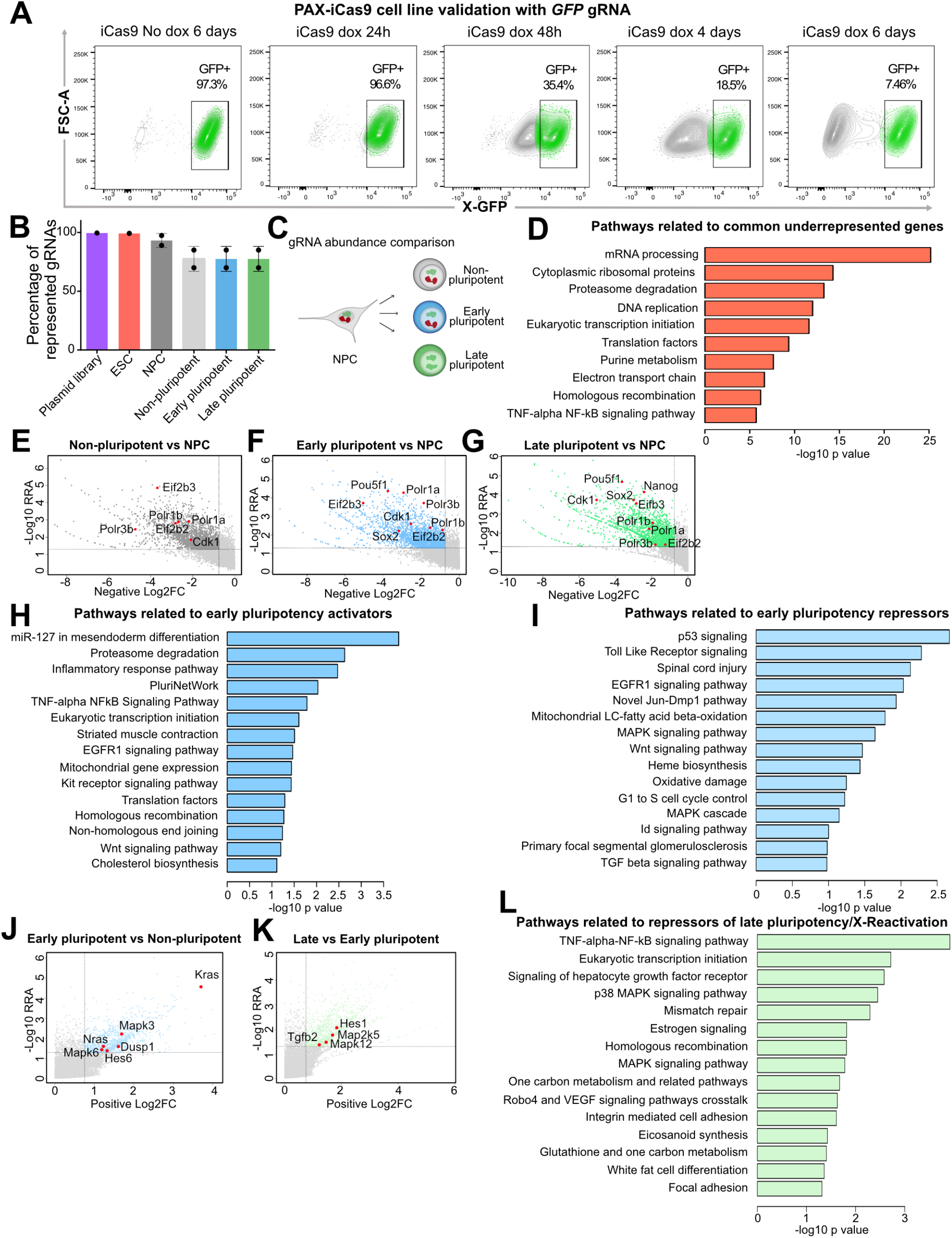
A genome-wide CRISPR knockout screen reveals molecular networks involved in reprogramming and X-chromosome reactivation. **Related to** Figure 1. (A) Validation of knockout efficiency by flow cytometry. Flow cytometry analysis during 6 days of doxycycline treatment in the X-GFP *iCas9* ESC line was done to measure the X-GFP percentage decay in cells containing a gRNA targeting the *GFP* gene. Gating shows the X-GFP+ population. (B) Percentage of gRNA representation in the plasmid library, infected ESCs and the 4 populations analyzed in two independent screening rounds: NPCs and day 10 reprogramming populations (non-pluripotent, early pluripotent, late pluripotent). Error bars represent SD. (**C**) gRNA abundance comparisons (related to D-G): NPCs to non-pluripotent, early pluripotent and late pluripotent populations. (**D**) Pathways related to common underrepresented genes (n=927 genes) in the three reprogramming populations compared to NPCs (WikiPathways Mouse 2019). For all comparisons, a RRA score < 0.05 and Log2FC < −0.75 (underrepresented) filtering was applied. (**E-G**) Representation of genes with negative Log2FC (underrepresented) vs −log10 RRA in the non-pluripotent (E), early pluripotent (F) and late pluripotent (G) populations compared to NPCs (RRA cutoff = 0.05, Log2FC cutoff = −0.75). (**H**) Pathways (WikiPathways Mouse 2019) related to underrepresented genes in the “early pluripotent vs non-pluripotent” comparison (activators of early pluripotency, n=1361 genes) (RRA score < 0.05 and Log2FC < −0.8 filtering was applied). (**I**) Pathways (WikiPathways Mouse 2019) related to overrepresented genes in the “early pluripotent vs non-pluripotent” comparison (repressors of early pluripotency, n=693 genes) (RRA score < 0.05 and Log2FC < −0.8 filtering was applied). (**J**) Representation of genes with positive Log2FC (overrepresented) vs −log10 RRA (RRA cutoff = 0.05, Log2FC cutoff = 0.75) in the “early pluripotent vs non-pluripotent” comparison (repressors of early pluripotency). (**K**) Representation of genes with positive Log2FC (overrepresented) vs −log10 RRA (RRA cutoff = 0.05, Log2FC cutoff = 0.75) in the “late pluripotent vs early pluripotent” comparison (repressors of late pluripotency, X-reactivation). (**L**) Pathways (WikiPathways Mouse 2019) related to overrepresented genes in the “late pluripotent vs early pluripotent” comparison (repressors of late pluripotency, X-reactivation, n=839 genes) (RRA score < 0.05 and Log2FC < −0.8 filtering was applied).

**Figure S2.**
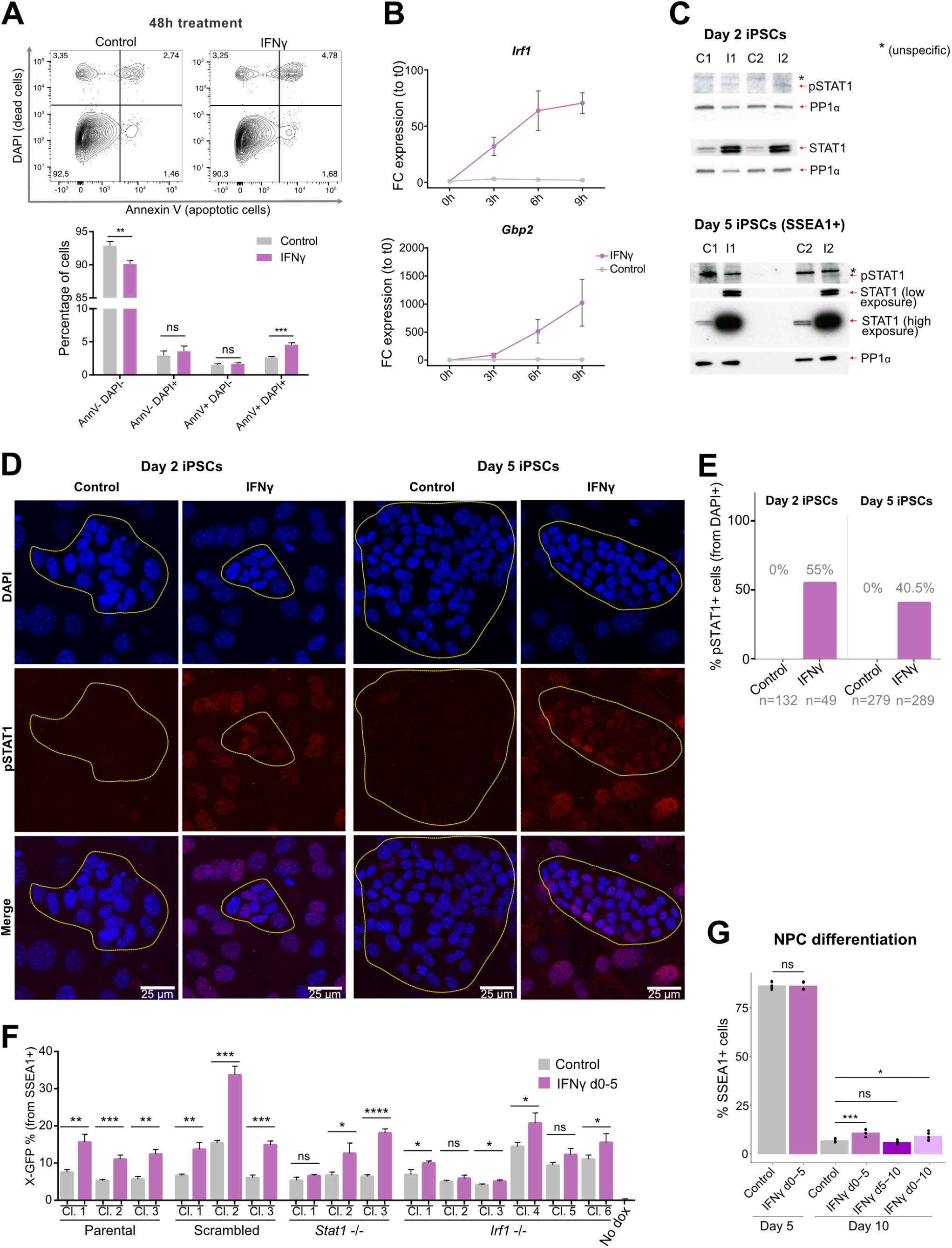
Interferon γ pathway activation during iPSC reprogramming. **Related to** Figure 2. (**A**) Analysis of apoptosis by annexin V and DAPI staining with flow cytometry after 48h of reprogramming induction +/− IFNγ treatment (n=3 technical replicates). (**B**) RT-qPCR on mRNA for *Irf1* and *Gbp2* expression at 0h, 3h, 6h and 9h from reprogramming induction +/− IFNγ treatment (relative to t0). Error bars represent SD (n=3 technical replicates). (**C**) Western blotting of STAT1 and pSTAT1 (Tyr701) on day 2 and day 5 iPSCs +/− IFNγ treatment (loading control: PP1α). (**D**) Immunofluorescence of pSTAT1 (Tyr701) on day 2 and day 5 iPSCs +/− IFNγ treatment. Scale bar = 25 µm. Outlines highlight colonies of cells undergoing reprogramming, characterized by smaller nuclei and tight aggregation. (**E**) Percentage of pSTAT1-positive cells from immunofluorescence in (D). Numbers of counted cells are indicated on the bottom of the graph. (F) (Related to Figure 2F-H). Flow cytometry quantification of total X-GFP percentages (from SSEA1+ cells) on day 7 of reprogramming for 3 clones from the parental cell line, 3 clones containing a scrambled gRNA, 3 *Stat1 −/−* clones and 6 *Irf1 −/−* clones, including three technical replicates for each clone, in IFNγ-treated cells and untreated controls. Statistics (unpaired t-tests): ns = non significant; * = p<0.05; ** = p<0.01, *** = p<0.001, **** = p<0.0001. Error bars represent SD. (**G**) (Related to Figure 2I-K). Quantification of SSEA1 percentage on days 5 and 10 of NPC differentiation by flow cytometry in control and IFNγ treatment conditions. Statistics (paired t-tests): ns = non significant; * = p<0.05; *** = p<0.001.

**Figure S3.**
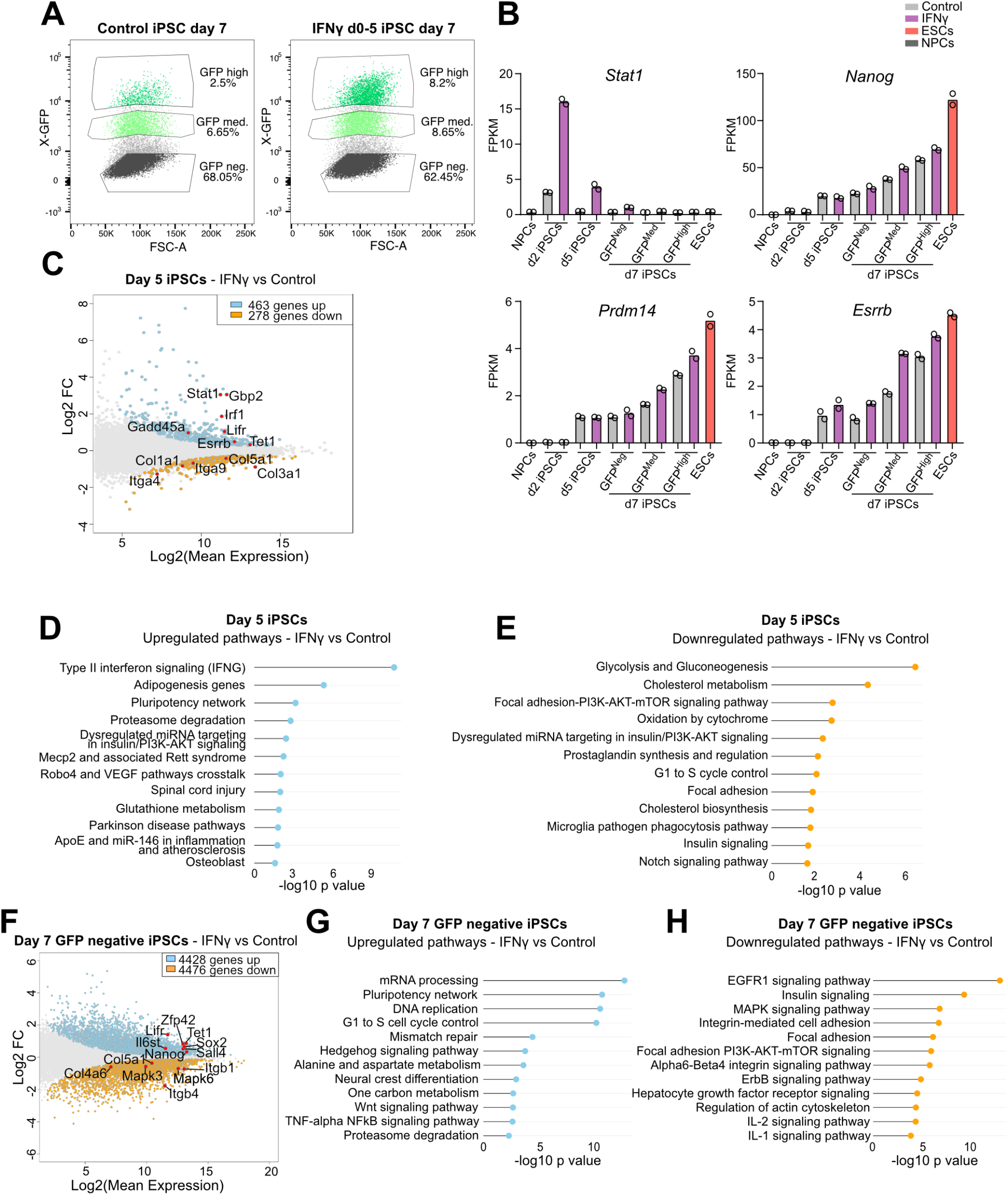
Transcriptomic analysis of interferon γ pathway activation during iPSC reprogramming. **Related to** Figure 3. (**A**) Flow cytometry plots of X-GFP expression (from SSEA1+ cells) in control and IFNγ-treated day 7 iPSCs. Gating shows sorted populations for RNA-sequencing. Average percentages between two independent reprogramming inductions are indicated for each population. (**B**) Expression (FPKM) of selected genes (*Stat1, Nanog, Prdm14* and *Esrrb*) in NPCs, ESCs, day 2, day 5 and day 7 iPSC populations +/− IFNγ treatment (two RNA-sequencing replicates shown). (**C**) MA plot displaying transcriptomic changes of IFNγ vs control d5 iPSCs (adjusted p value = 0.1). Upregulated genes are highlighted in light blue, downregulated genes are highlighted in orange. Selected genes are shown with points in red. (**D, E**) Upregulated (D) and downregulated (E) pathways of IFNγ vs control d5 iPSCs (WikiPathways Mouse 2019) (adjusted p value = 0.1). (**F**) MA plot displaying transcriptomic changes of IFNγ vs control d7 X-GFP negative iPSCs (adjusted p value threshold = 0.1). Upregulated genes are highlighted in light blue, downregulated genes are highlighted in orange. Selected genes are shown with points in red. (**G, H**) Upregulated (G) and downregulated (H) pathways (WikiPathways Mouse 2019) IFNγ vs control d7 X-GFP negative iPSCs (WikiPathways Mouse 2019) (adjusted p value = 0.1).

**Figure S4.**
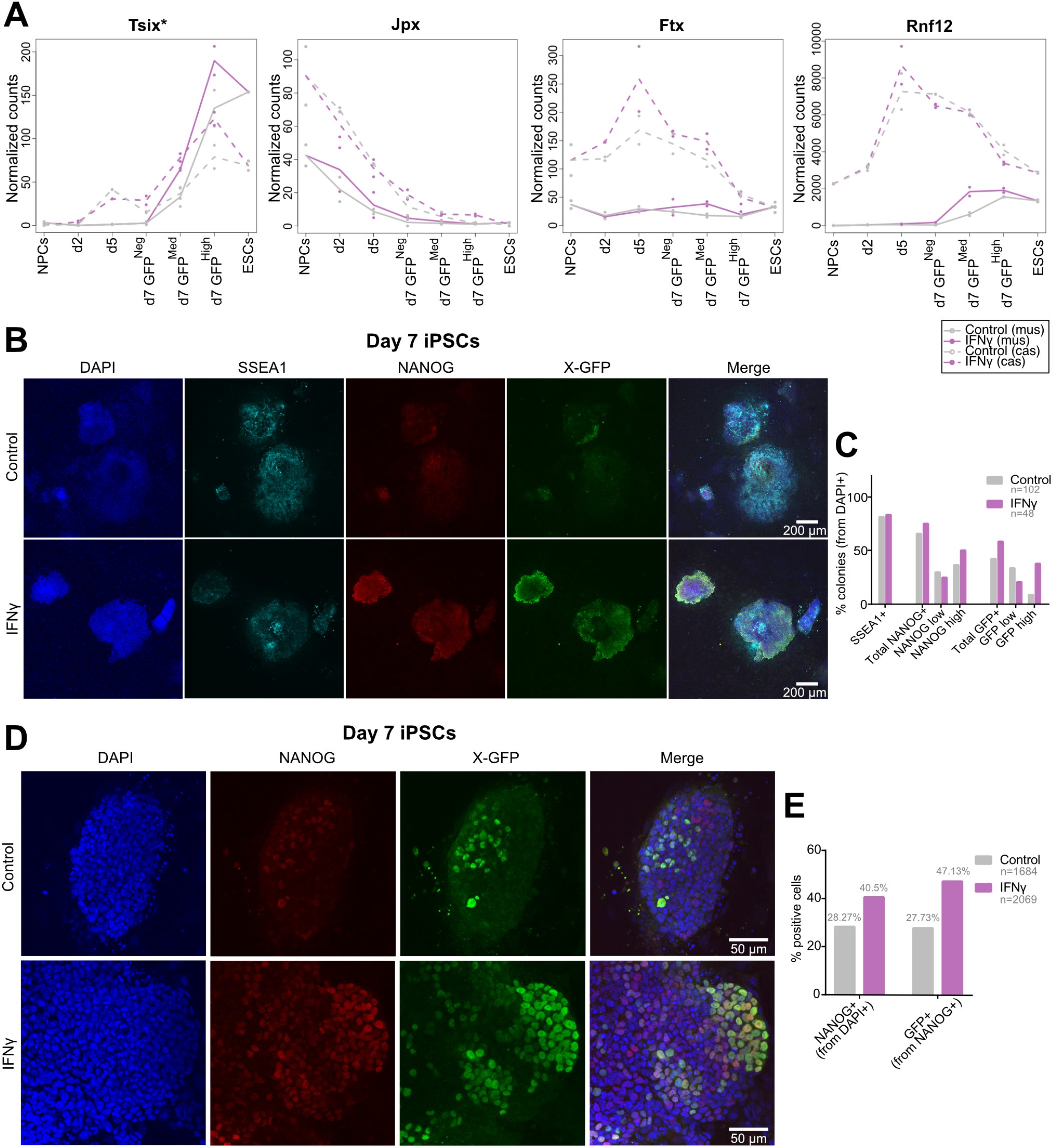
Increased expression of NANOG and X-GFP in iPSC colonies upon early interferon γ treatment. **Related to** Figure 4. (**A**) Expression (normalized counts) of genes of the X-inactivation center (*Tsix*, *Jpx*, *Ftx*, *Rnf12*) from X mus and X cas on NPCs, ESCs, day 2, day 5 and day 7 iPSC populations +/− IFNγ treatment (two RNA-sequencing replicates shown). The * at *Tsix* indicates that the gene contains a truncation on the X-mus (*100*) and therefore cannot regulate *Xist* expression *in cis*. (**B**) Immunofluorescence (low magnification, 4×) for SSEA1, NANOG and X-GFP (active X chromosome) of day 7 reprogramming colonies +/− IFNγ treatment. Scale bar = 200 µm. (**C**) Percentages of SSEA1+, NANOG+ (low/high) and X-GFP+ (low/high) colonies from immunofluorescence in (B). The number (n) of counted colonies is indicated in the graph. NANOG+ or X-GFP+ colonies were scored as low or high if approximately less or more than half of the cells in the colony were positive for these markers, respectively. (**D**) Immunofluorescence (high magnification, 63x) for NANOG and X-GFP (active X chromosome) of day 7 reprogramming colonies +/− IFNγ treatment. Scale bar = 50 µm. (**E**) Percentages of NANOG+ and X-GFP+ (from NANOG+) cells from immunofluorescence in (D). The number (n) of counted cells is indicated in the graph.

**Figure S5.**
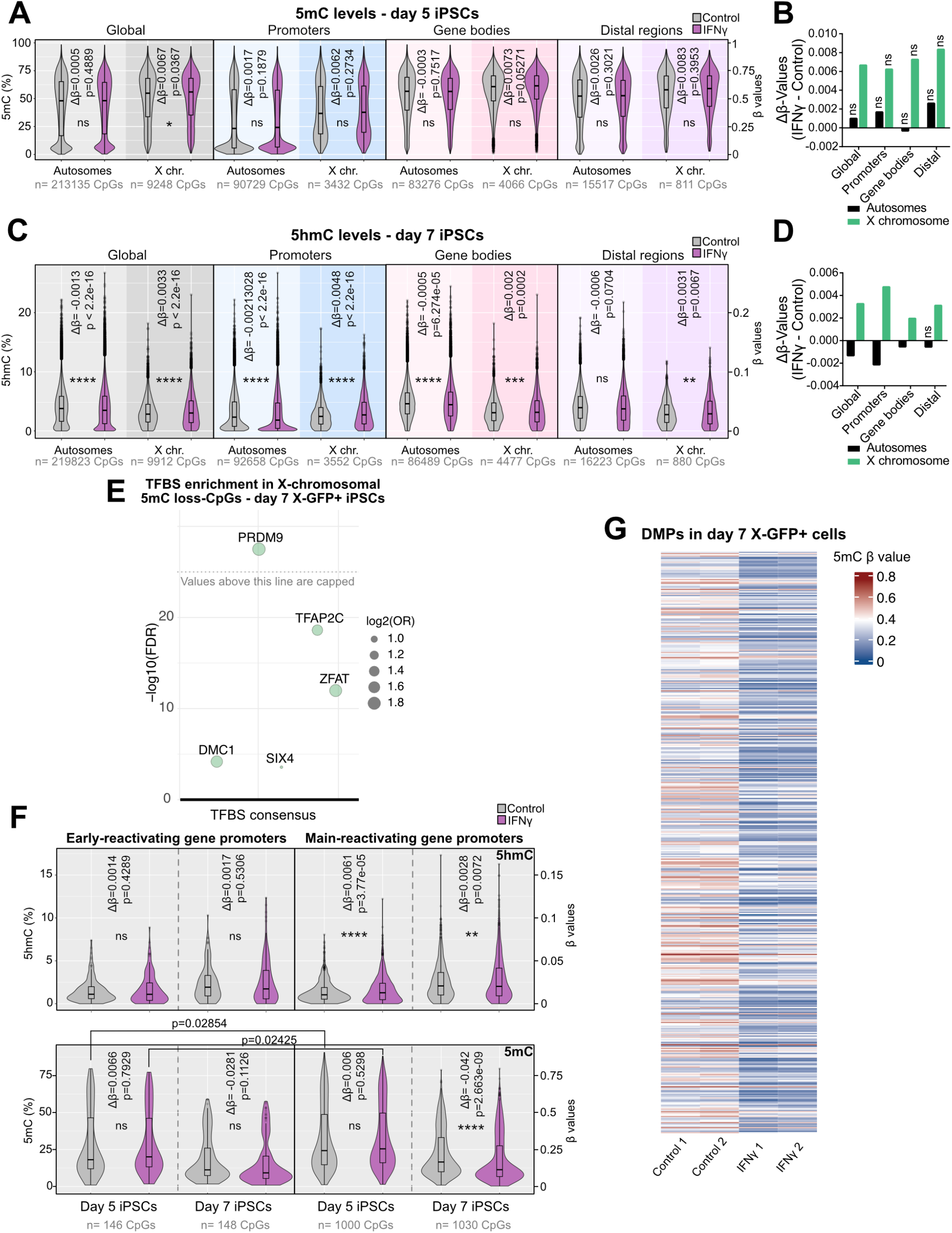
Interferon γ treatment promotes active DNA demethylation in cells undergoing reprogramming. **Related to** Figure 5. (A) Analysis of 5mC levels (β-values) of CpGs in autosomes and X chromosome in day 5 iPSCs for control and IFNγ conditions, globally and divided by genomic distribution: promoters (<= 1kb from TSS), gene bodies and distal regions (number (n) of detected CpGs from each category is indicated on the bottom of the graphs). Δβ-values (mean β-value IFNγ - mean β-value control) and p values (comparison IFNγ vs control) are shown in the graphs. Statistics: unpaired t-tests. (B) Δβ-values (mean β-value IFNγ - mean β-value control) for 5mC in day 5 iPSCs for each genomic region (global, promoters, gene bodies and distal regions) in autosomes and X chromosomes (corresponding to analysis in (A)). Bars marked with “ns” correspond to non-significant changes from analysis in (A). (**C**) Analysis of 5hmC levels (β-values) of CpGs in autosomes and X chromosome in day 7 X-GFP+ iPSCs for control and IFNγ conditions, globally and divided by genomic distribution: promoters (<= 1kb from TSS), gene bodies and distal regions (number (n) of detected CpGs from each category is indicated on the bottom of the graphs). Δβ-values (mean β-value IFNγ - mean β-value control) and p values (comparison IFNγ vs control) are shown in the graphs. Statistics: unpaired t-tests. (**D**) Δβ-values (mean β-value IFNγ - mean β- value control) for 5hmC in day 7 X-GFP+ iPSCs for each genomic region (global, promoters, gene bodies and distal regions) in autosomes and X chromosomes (corresponding to analysis in (C)). Bars marked with “ns” correspond to non-significant changes from analysis in (C). (**E**) Transcription factor binding site (TFBS) enrichment analysis on differentially methylated X-chromosomal CpGs (DMPs, logFC<(−0.1), p<0.01, n=468 CpGs) which lose methylation upon IFNγ treatment compared to control in day 7 X-GFP+ iPSCs. Analysis was performed with Sesame R package. (**F**) Analysis of 5mC and 5hmC levels (β-values) of CpGs in early and main X-reactivating gene promoters at day 5 and day 7 X-GFP+ iPSCs for control and IFNγ conditions (gene lists were obtained from Bauer et al, 2021). Number (n) of detected CpGs for each category and time point is indicated on the bottom of the graphs. Δβ-values (mean β-value IFNγ - mean β-value control) and p values (comparison IFNγ vs control) are shown in the graphs. Statistics: unpaired t-tests. (**G**) Heatmap showing 5mC levels (β-values) of all X-chromosomal differentially methylated CpGs (n=470 DMPs, logFC cutoff = +/−0.1, p<0.01) sorted by chromosome position.

## Notes

### Competing Interest Statement

The authors have declared no competing interest.

### Summary of Updates

The spelling of interferon gamma was corrected in the title and the abstract in order to use the greek letter for gamma.

